# The conserved Arabidopsis PM19L1 protein functions as an osmosensor and regulator of dormancy and germination

**DOI:** 10.1101/2020.08.10.244889

**Authors:** Ross D. Alexander, Pablo Castillejo-Pons, Nina Melzer, Omar Alsaif, Vivien I. Strotmann, Yvonne Stahl, Madeleine Seale, Peter C. Morris

## Abstract

How plants perceive water, especially during the critical stages of seed formation and germination, is key to their survival. During development and germination, seeds undergo large changes in water content, down to around 10% during maturation and up to 90% again within 24 hours of germination. We find the evolutionary conserved Arabidopsis plasma membrane protein PM19L1 to be an osmosensor, regulating dormancy and germination under osmotic stress. The PM19L1 protein structurally resembles the yeast osmosensor Sho1, with four transmembrane domains, and *PM19L1* complements the osmosensitive *sho1* mutant. Arabidopsis *pm19l1* mutants have enhanced dormancy and reduced germination under salt and osmotic stress, and enhanced ABA levels. In a striking parallel to osmosensing in yeast, signalling downstream of PM19L1 involves a MAP kinase signal transduction pathway. *PM19L1* is a positive regulator of *ABI3*, which promotes the late maturation of the seed, and negatively regulates the *ABI4* and *ABI5* dormancy- regulating transcription factors. These results have implications for the study of dormancy, drought, and salinity tolerance in crops, and may provide an insight into evolutionary adaptation of plants to a terrestrial environment.

## Introduction

Abiotic stress is a major constraint on agricultural productivity. Drought and salinity are particularly problematic and are exacerbated by anthropogenic climate change. Many organisms have mechanisms for perceiving and thus adapting to environmental osmotic stress but perception mechanisms in plants have remained elusive (Haswell and Verslues, 2015; Nongpiur et al., 2020). The AWPM-19 (ABA-induced wheat plasma membrane polypeptide) protein was first identified in the plasma membrane fraction of abscisic acid (ABA) treated wheat cells (Koike et al., 1997), and subsequently in dormant embryos of barley (Ranford et al., 2002). AWPM-19 homologues have been associated with preharvest sprouting, seed dormancy and drought tolerance in a number of different species (Barrero et al., 2015; Chen et al., 2015; Yao et al., 2018; Barrero et al., 2019), and expression of AWPM-19-like genes is enhanced in rice and Arabidopsis by osmotic stress, drought, salt, ABA and cold (Chen et al., 2015; Yao et al., 2018; Barrero et al., 2019). In the model plant *Arabidopsis thaliana*, AWPM-19 proteins are encoded for by a 4-member gene family (*PM19L1, PM19L2, PM19L3, PM19L4*) (Barrero et al., 2019). In this work we show that the AWPM-19 protein encoded by *PM19L1* (AT1G04560) functions as an osmosensor and can regulate dormancy and germination through the control of key transcription factors ABI3 (ABA-Insensitive 3) (Koornneef et al., 1984), ABI4 (ABA Insensitive 4) and ABI5 (ABA Insensitive 5) (Finkelstein, 1994)). In the context of the seed, ABI3 is required for seed maturation and imposing a transition between embryo maturation and seedling development, whereas ABI4 and ABI5 help regulate post germination arrest of seedling growth through enhanced sensitivity to ABA under stress conditions, as well as having a role in seed maturation together with ABI3 (reviewed, Finkelstein et al., 2002; Ali et al., 2022).

## Results

Analysis of the structure of the deduced PM19L1 protein sequence suggests the presence of four transmembrane domains and a variable C-terminal tail (Fig. 1A). The amino acid sequences in the four predicted transmembrane domains and in the intracellular loop between helices two and three are highly conserved in all land plants examined, including bryophytes. This shows the evolutionary antiquity (approximately 470 million years) of this protein (Fig. 1B). Homologues were not found in any other organisms including algae (for example the Zygnematophyceae, the closest living relative of land plants (Cheng et al., 2019). These analyses imply that the transmembrane domains are not simply structural but have an important and conserved function. In contrast, the predicted extracellular loops and C-terminus are not conserved and are variable in length. We confirmed the plasma membrane localisation of PM19L1 protein (Koike et al., 1997; Chen et al., 2015; Yao et al., 2018) by confocal imaging of cells from transgenic Arabidopsis seedlings expressing free GFP or a PM19L1-GFP fusion driven by the CaMV 35S promoter. PM19L1-GFP exhibited colocalization with FM4-64 at the plasma membrane (Fig. 1C).

**Figure 1.**
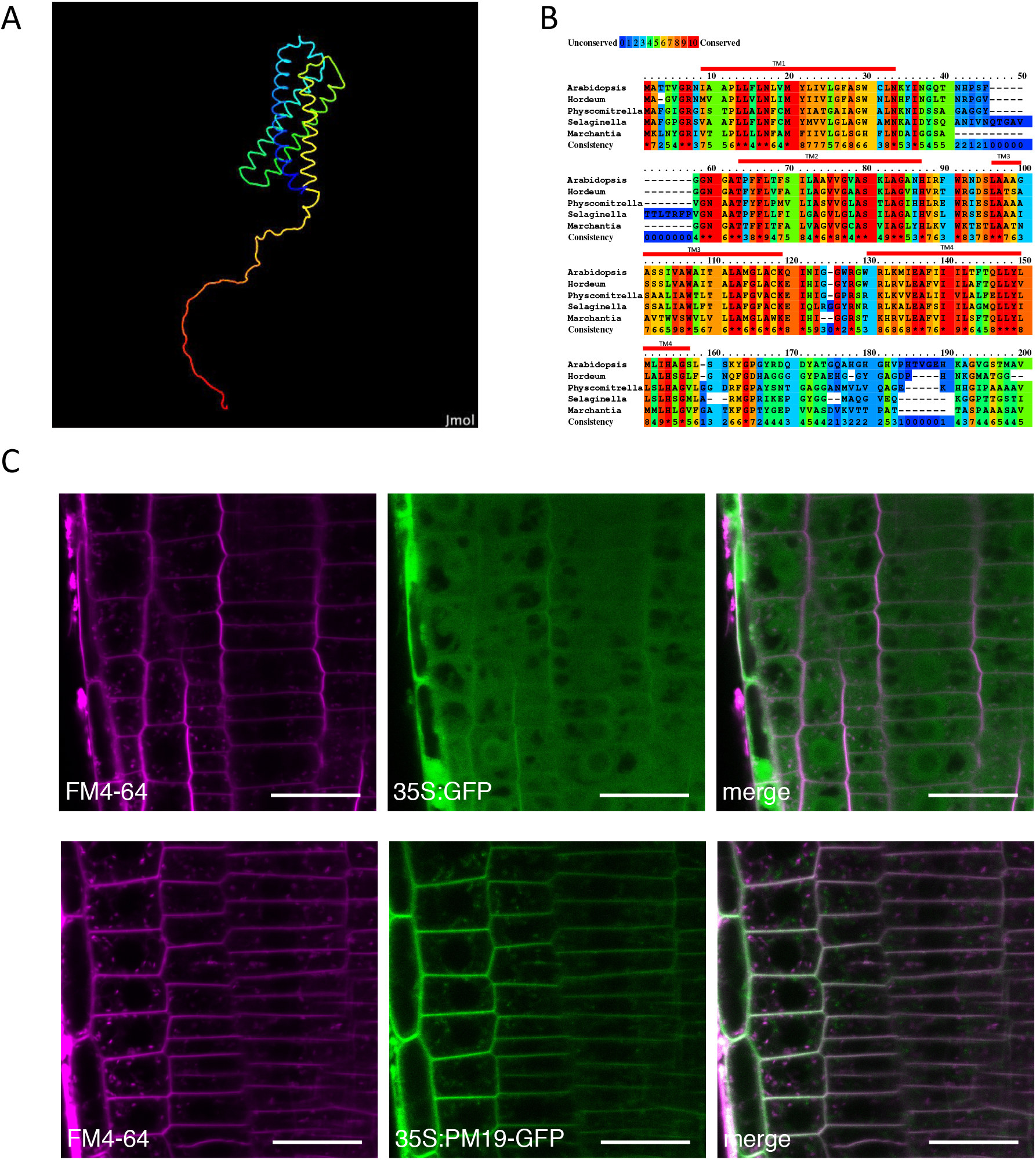
PM19L1 has four transmembrane domains and is highly conserved through terrestrial plant evolution. (A) Predicted secondary structure of PM19L1 protein carried out with AlphaFold (Jumper *et al*., 2021; Varadi *et al*., 2022) and visualised with JMol (Jmol: an open-source Java viewer for chemical structures in 3D. http://www.jmol.org/), showing 4 transmembrane alpha helixes and a C-terminal tail (N terminal coloured blue, C terminal red). (B) Alignment analysis of PM19L1 proteins from representative members of land plants, including bryophytes, vascular plants, and seed plants. Red bars indicate position of predicted transmembrane domains. Uniprot sequences *Arabidopsis thaliana* 023029 (the encoded protein from *PM19L1*), *Hordeum vulgare* A0A287T362, *Physcomitrella patens* A9SMF8, *Selaginella moellendorffii* D8S1W8 and *Marchantia polymorpha* A0A2R6WWV5 were aligned using Praline multiple sequence alignment (http://ibi.vu.nl/programs/pralinewww/). (C) Intracellular localisation of free GFP (top row) and PM19L1-GFP (bottom row) (scale bar 20 µm) in transgenic Arabidopsis seedling roots as visualised by confocal microscopy. Cells were counterstained with FM4-64 (magenta) to visualise the plasma membrane.

Transgenic Arabidopsis plants bearing the *PM19L1* promoter sequence fused to the UidA marker gene show high levels of GUS activity in the germinating seedling, but not in vegetative tissues, other than in root tips (Fig. 2A, B). Transcripts of *PM19L-1* mRNA are highest during seed development and early germination but are also observed in all other tissues at lower levels. Expression of the three other gene family members was lower than *PM19L1* in seeds but was seen in all tissues including stems and roots (Fig. 2C, D). Polyethylene glycol (PEG), salts and ABA all upregulated *PM19L1* expression in 14-day old seedlings, with PEG producing a particularly strong effect. Little effect was seen for the other gene family members (Fig. 2E).

**Figure 2.**
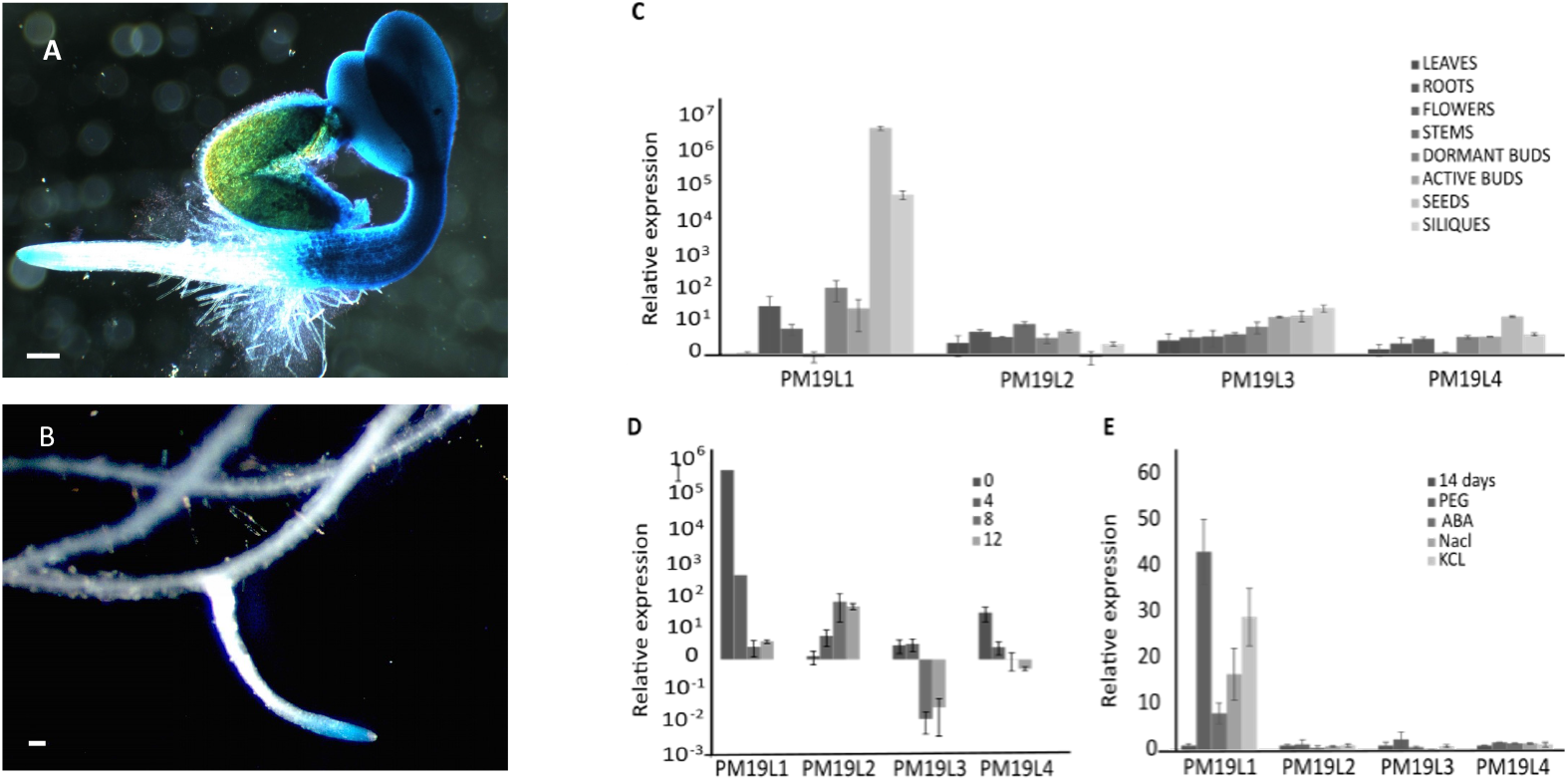
*PM19L1* is highly expressed during germination and in root tips. Tissue-specific localisation of *PM19L1* gene expression as visualised by staining seedlings bearing the *pPM19L1:UidA* construct for GUS activity, showing high levels of expression in germinating seedlings (A) and in lateral root tips (B) of 14 day old seedlings (scale bars 100 µm). Q-PCR analysis of expression patterns for Arabidopsis *PM19L* gene family members *PM19L1* (AT1G04560), *PM19L2* (AT5G46530), *PM19L3* (AT1G29520) and *PM19L4* (AT5G18970) in (C) different organs, (D) during germination (bars show days of germination) and (E) in seedlings in response to osmotic stresses (15% PEG 6000, 200 mM NaCl, 200 mM KCl) and 25 µM ABA; 14 days indicates untreated control plants. Bars show standard deviation.

An analysis of the phylogenetics for AWPM-19 proteins in representatives of all land plant families shows that the Arabidopsis PM19L1 protein clusters with other monocot and eudicot AWPM-19 representatives, and most of the AWPM-19 representatives from the bryophytes. In contrast, PM19L2-4 are found in a separate clade and are more divergent from bryophyte sequences (Fig. S1, Table S1,2).

Given the predominantly seed and seedling-specific expression pattern of *PM19L1*, and the induction of *PM19L1* gene expression by osmotica and salts, we suspected that PM19L1 may play a role in germination. We analysed a T-DNA insertion mutant for *PM19L1*, (which did not express detectable mRNA for *PM19L1,* Fig. S2), for germination-specific traits. The *pm19l1* mutant was found to be more sensitive than the wildtype to the presence of salt or sorbitol in the germination medium. (Fig. 3A, B). The greater inhibitory effect of NaCl compared to equimolar sorbitol is characteristic of the cytotoxic effect of sodium. The decreased germination on NaCl or sorbitol was rescued by complementing the mutant line either with a constitutive promoter driving expression of a *PM19L1-GFP* fusion (*pCaMV 35S*:*PM19L1-GFP*), or by the native *PM19L1* promoter and gene (*pPM19L1:PM19L1*). The expression pattern and intracellular location of *PM19L1*, together with the germination phenotype of the *pm19l1* mutant strongly suggests a protein function associated with perception of osmotic stress.

**Figure 3.**
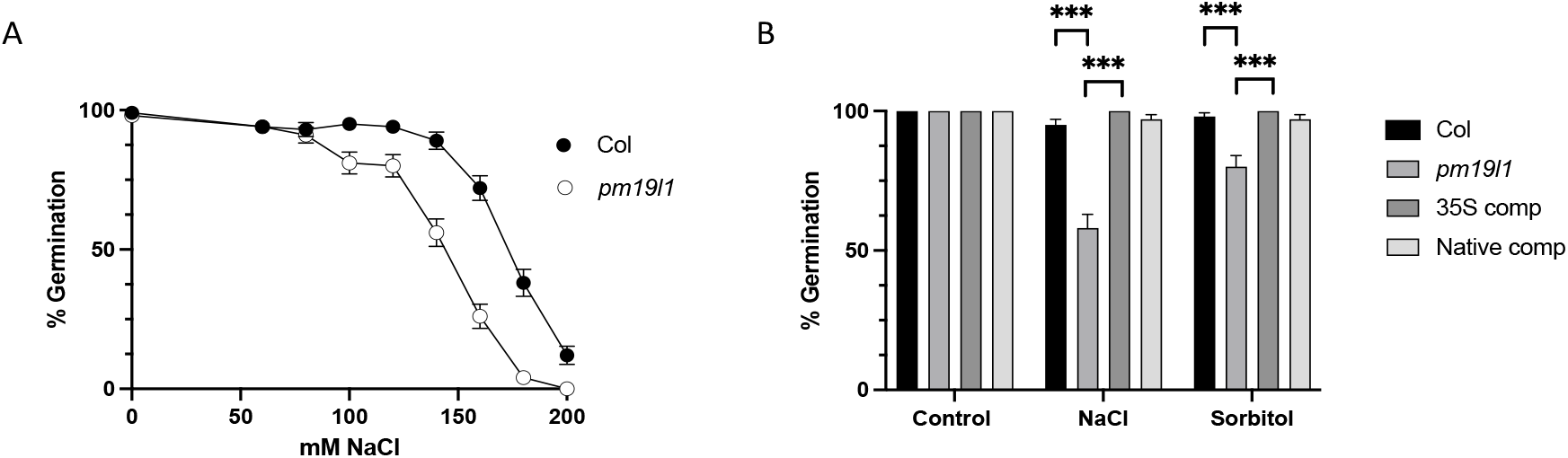
The *pm19l1* mutant is sensitive to salt and osmotic stress during germination. (A) germination of Columbia wildtype and *pm19l1* mutant seeds on media with varying concentrations of NaCl. (B) Germination of Columbia wildtype, *pm19l1*and lines complemented with *pCaMV 35S:PM19L1-GFP* or the native promoter (*pPM19L1:PM19L1*) on control media and media containing 100 mM NaCl or 200 mM sorbitol. Bars show standard deviation (based on binomial distribution, 100 seeds). Statistically significant differences, using Fisher’s exact test followed by a Bonferroni-Šídák post-hoc analysis are indicated with *** showing P ≤ 0.001.

We explored the possible role of the PM19L1 protein by genetic complementation of different yeast mutants that have phenotypes associated with salt and osmotic stress. Sho1 is an osmosensor that permits growth of yeast on media of high osmolarity by stimulating the Hog1 signalling pathway that controls the accumulation of the osmolyte glycerol (Maeda et al., 1995; Saito and Posas, 2012; Tatebayashi et al., 2020). The *sho1* mutant was fully complemented when PM19L1 was expressed, permitting growth on up to 500 mM NaCl or sorbitol. The deletion of the sequence encoding the variable C-terminus of PM19L1 had no impact on the ability of PM19L1 to complement *sho1* (Fig. 4A, B). Sho1, like PM19L1, is a four transmembrane domain plasma membrane protein (Fig. S3) with an extended C terminal tail but has no protein sequence homology with the AWPM-19 proteins. The ability of an unrelated Arabidopsis four-transmembrane domain plasma membrane protein, tetraspanin 3, was tested for its ability to complement *sho1*, but it failed to do so (Fig. S4). This shows that the PM19L1 protein is specific in its ability to function as an osmosensor and suggests a role for PM19L1 in the control of germination through sensing of the osmotic environment. Other potential mechanisms for the role of PM19L1 were also explored using yeast mutants. No complementation was seen for the potassium influx transporter mutants *trk1* and *trk2* (Bertl et al., 2003) (Fig. S5) nor the sodium efflux mutant *ena1* (Haro et al., 1991) (Fig. S6). The yeast glycerol biosynthetic mutant *gdp1/gdp2* (Pettersson et al., 2006) was also not complemented, indicating that PM19L1 does not have an aquaporin-like function (Fig. S7).

**Figure 4.**
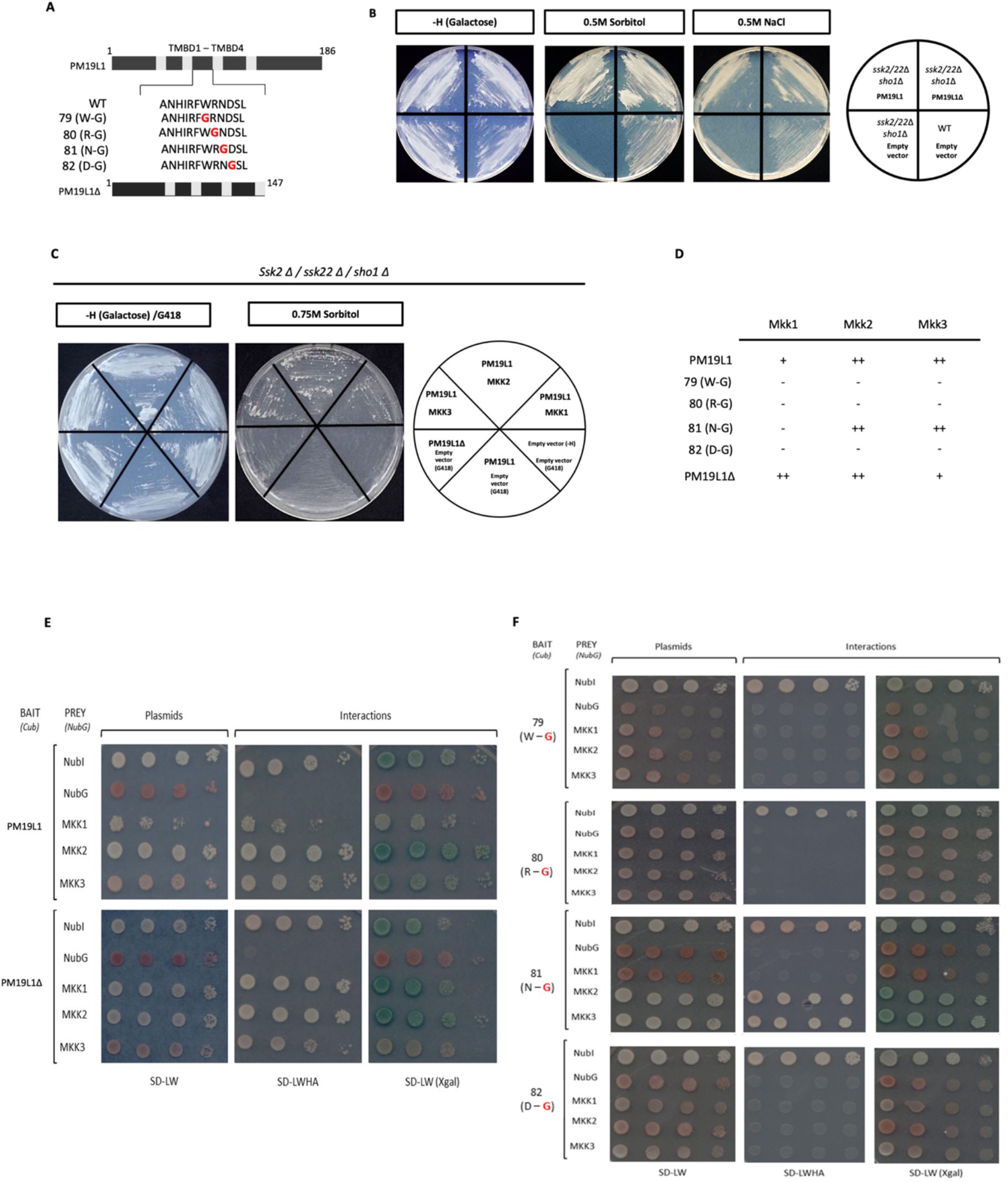
Complementation of the *sho1* mutant in yeast by expression of *PM19L1* and interaction with MAP kinases kinase proteins. (A) shows the constructs tested, expression of full-length *PM19L1* (plasmid pPMRA5), and truncated *PM19L1* with the variable C terminus removed (*PM19L1Δ*)(plasmid pPMRA6). (B) shows growth of wildtype, *skk1/skk2/sho1* mutant, *skk1/skk2/sho1* transformed with *PM19L1*, and with *PM19L1Δ* on basal media, 0.5 M sorbitol and 0.5 M NaCl. (C) co-expression of Arabidopsis MAP kinase kinase (*MKK1*, *MKK2* and *MKK3*) enhances the ability of *PM19L1* to complement *sho1*, resulting in growth on up to 0.75 M sorbitol. (D) is a summary of split ubiquitin analysis of PM19L1 interactions with Arabidopsis MKK proteins. (E) Split ubiquitin assays showing interactions between PM19L1 or PM19L1*Δ* with Arabidopsis MKK proteins. NubI is a positive control, NubG a negative control, SD-LW medium selects for presence of bait and prey proteins, SD-LWHA shows growth on media minus histidine and adenine, and SD-LW (Xgal) shows activity of LacZ reporter gene (blue colour). (F) Split ubiquitin assays showing interactions between PM19L1 or PM19L1*Δ* with Arabidopsis MKK proteins after site- directed mutagenesis to glycine of conserved (79 W-G, 80 R-G, and 82 D-G) or non- conserved (81 D-G) amino acids in the intracellular loop of PM19L1.

The yeast osmosensing pathway initiates from two osmosensors, the histidine kinase Sln1 and the four-transmembrane domain protein Sho1. Downstream from both osmosensors is a kinase signal transduction pathway that is activated under conditions of high osmolarity, with Sln1 activating Ssk2 and Ssk22 (two redundant MAP kinase kinase kinases, MKKK), and the MKKK Ste11 being activated by Sho1. Both Ssk2/Ssk22 and Ste11 will in turn activate the MAP kinase kinase (MKK) Pbs2 (which physically interacts with Sho1), and then Pbs2 activates the Map kinase Hog1, resulting in the accumulation of glycerol as an osmolyte and the expression of aquaporins to permit water ingress (O’Rourke et al., 2002; Tatebayashi et al., 2015). The *pbs2* mutant is compromised by osmotic stress, however growth of *pbs2* on media of high osmolarity is restored by expression of Arabidopsis MKK2 (Ichimura et al., 1998). If Arabidopsis MAP kinase signalling is also involved in plant osmosensing downstream of PM19L1, then co-expression of Arabidopsis MKK genes together with PM19L1 in the yeast *sho1* mutant background might be expected to enhance yeast growth under osmotic stress. This was found to be the case for *AtMKK1*, *AtMKK2* and *AtMKK3*. All three of these Arabidopsis MKK genes have previously been reported to be associated with germination stress, with mutants more sensitive to salt, although we have not been able to confirm this for AtMKK1 (Fig. S8) (Teige et al., 2004; Hwa and Yang, 2008; Conroy et al., 2013). When introduced into the *sho1* mutant, co-expression of the Arabidopsis MKKs permitted growth on up to 0.75 M sorbitol, whereas PM19L1 on its own only permitted growth on up to 0.5 M sorbitol (Fig. 4C), suggesting that Arabidopsis MKK proteins may be associated with downstream signalling from PM19L1, in a similar manner to Pbs2 and Sho1. This association was corroborated by a split ubiquitin analysis in which PM19L1 (and C-terminally truncated PM19L1) was found to interact with AtMKK1 (weak interaction), AtMKK2 and AtMKK3 (which both showed a stronger interaction (Fig. 4E). MKKs are cytosolic and thus might be expected to interact with the intracellular domains of PM19L1. Site directed mutagenesis of conserved amino acids within the predicted intracellular loop between transmembrane domains 2 and 3 (79 W-G; 80 R-G, 82 D-G) abolished the interaction, however this was not the case (apart for MKK1) when a non-conserved amino acid (81 N-G) in the same loop was changed (Fig. 4D, F). Complementation of *sho1* by *PM19L1* expression was abolished by mutagenesis of the conserved amino acids (Fig. S9).

The interaction of PM19L1 with MKK proteins was confirmed by carrying out pull-down assays with immobilised recombinant MBP-MKK1, MBP-MKK2 and MBP-MKK3 proteins incubated with extracts from plants expressing free GFP or PM19L1-GFP. MBP-MKK2 and MBP-MKK3, but not MBP-MKK1, specifically interacted with PM19L1-GFP in this assay (Fig. 5), which accords with the weak MKK1-PM19L1 interaction seen in the split ubiquitin assay.

**Figure 5.**
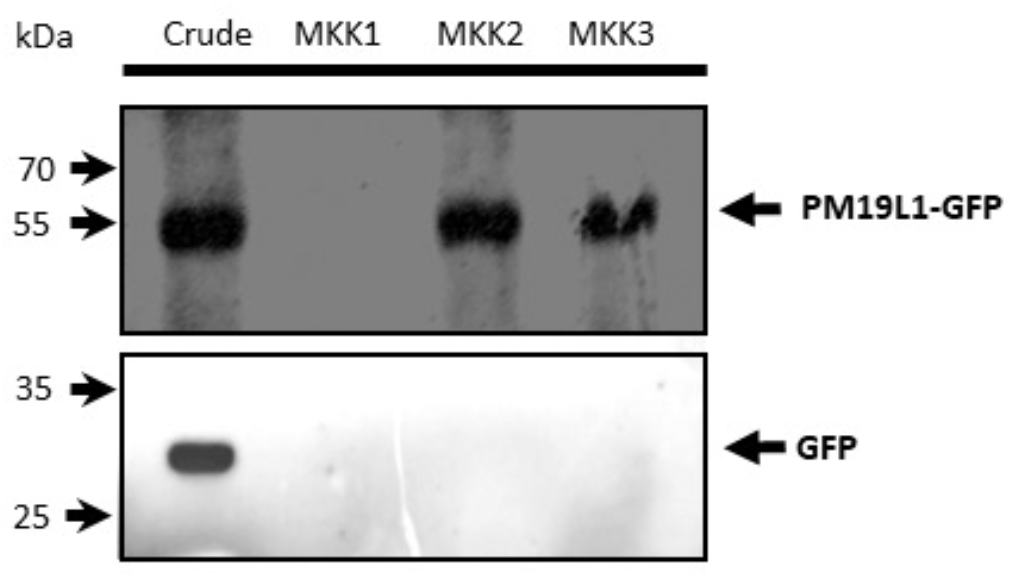
Pulldown assay showing interaction between MKK proteins and PM19L1. Immobilised MBP-MKK was incubated with extracts from plants expressing PM19L1-GFP or free GFP. Bound proteins and crude extract were fractionated by SDS-PAGE followed by immunoassay for GFP. MBP-MKK2 and MBP-MKK3 bind to PM19L1-GFP but not to free GFP. Top panel, PM19L1-GFP, lower panel, free GFP.

The physiology of *pm19l1* seeds and seedlings was further investigated to gain an insight into the mechanism of action of this osmosensor in regulating germination. Levels of sodium and potassium ions in seedlings of the *pm19l1* mutant did not differ from the wildtype when grown on 60 mM sodium or potassium salts (Fig. S10), evidence against the possibility that the *PM19L1* gene encodes a potassium or sodium transporter. Osmolyte accumulation such as proline and total soluble sugars in salt-stressed seedlings were no different in wildtype and *pm19l1* seedlings (Fig. S11A, B), making it unlikely that PM19L1 plays a role in adaptation to osmotic stress.

Experiments using inhibitors of ABA and GA synthesis point towards an involvement of ABA in the *pm19l1* phenotype for salt sensitivity. Inclusion of 150 µM sodium tungstate, an inhibitor of ABA biosynthesis (Lee and Milborrow, 1997), partially rescued *pm19l1* germination inhibition by 150 mM NaCl (Fig. 6A). Gibberellins are plant hormones that promote germination and often act in opposition to ABA. Inclusion of gibberellic acid in the germination medium enhanced germination of *pm19l1* on 100 mM NaCl (Fig. 6B). Biosynthesis of gibberellin can be inhibited by paclobutrazol (Dalziel and Lawrence, 1984). Inclusion of paclobutrazol in the germination medium inhibited germination with a greater impact on *pm19l1*, suggesting that the *pm19l1* mutant requires more GA synthesis for germination than does the wildtype (Fig. 6C). Endogenous levels of ABA were measured in after-ripened seeds, the *pm19l1* mutant was found to have around four times the level of ABA as wildtype seeds (Fig. 6D). Transcript levels of *ABI3*, *ABI4* and *ABI5* were measured in after-ripened dry seeds of *pm19l1*, and in *mkk2* and *mkk3* mutant seeds. Relative to levels in wildtype seeds, for all the mutant lines the *ABI3* levels were reduced between 10 and 100- fold, and both *ABI4* and *ABI5* transcripts were enhanced some 5 to 10-fold (Fig. 6E). Genes for ABA and gibberellin metabolism were also investigated (Fig. 6F). NCED6 is a 9-cis-epoxycarotenoid dioxygenase involved in the early steps of ABA biosynthesis (Lefebvre et al., 2006). *NCED6* transcripts were elevated relative to wildtype in *pm19l1*, *mkk2* and *mkk3* seeds, indicative of enhance ABA synthesis. CYP707A1 is an ABA hydrolase (Okamoto et al., 2006), and transcripts for the corresponding gene were reduced in *pm19l1* and *mkk3*. GA20OX1 catalyses conversion of GA12 to GA9 (Phillips et al., 1995), and transcripts for this gene were downregulated in *pm19l1* and *mkk3*. This suggests that for seeds of these mutants, ABA biosynthesis is favoured whilst gibberellin synthesis is reduced.

**Figure 6.**
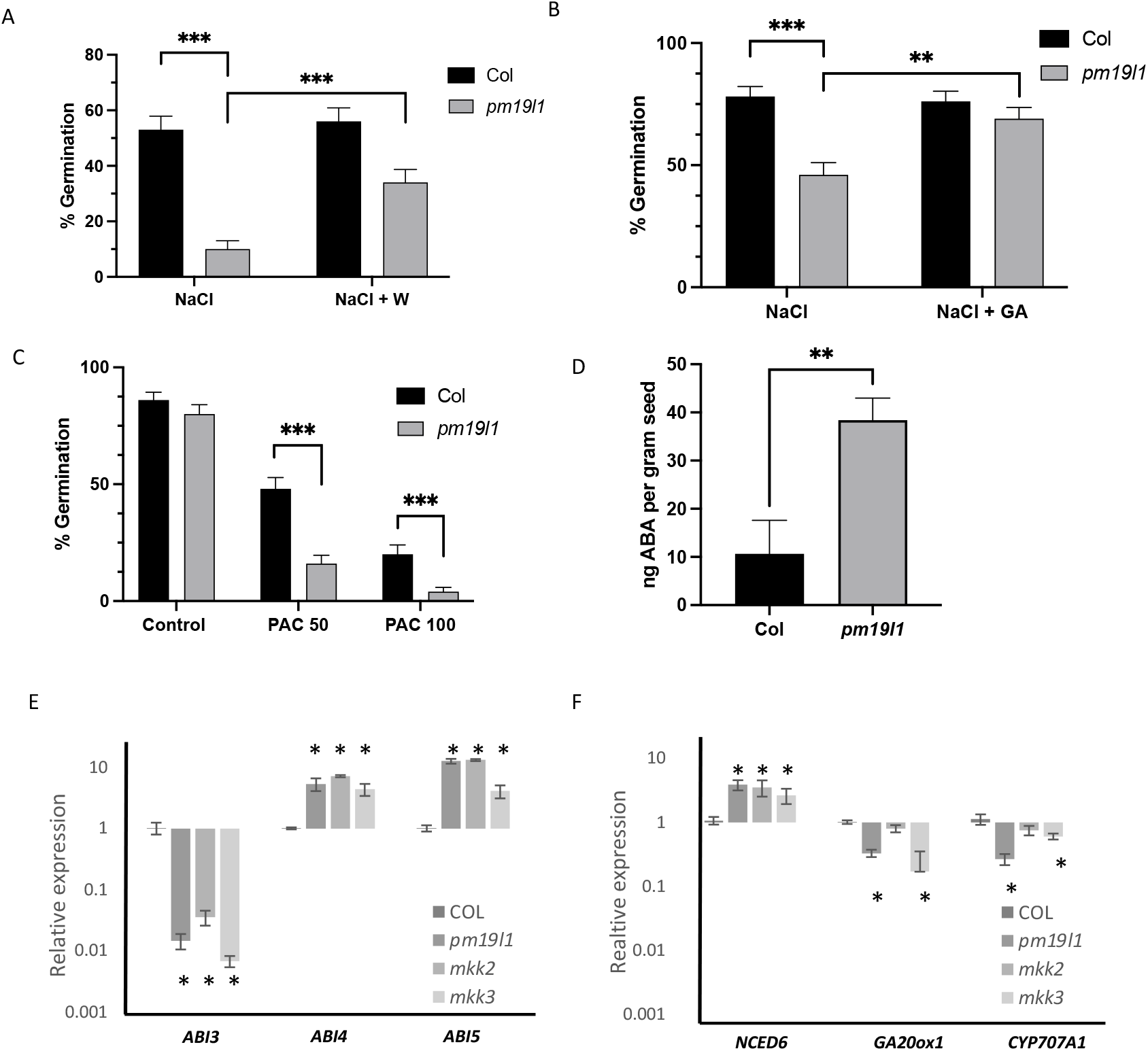
Role of abscisic acid in salt sensitivity. (A) After-ripened seeds were germinated for 7 days in media containing 150 mM NaCl with and without 150 µM of the ABA synthesis inhibitor sodium tungstate (W); seeds of *pm19l1* are partially rescued from salt inhibition by growth on tungstate. (B) Seeds were germinated in media containing 100 mM NaCl with and without 1 µM gibberellin (GA); seeds of *pm19l1* had enhanced germination in the presence of exogenous gibberellin. (C) Seeds were germinated in the presence of different concentrations (10-100 µM) of the GA synthesis inhibitor paclobutrazol (PAC); *pm19l1* seeds were more strongly inhibited from germination than the wildtype. Bars show standard deviation (based on binomial distribution, 100 seeds). Statistically significant differences, using Fisher’s exact test followed by a Bonferroni-Šídák post-hoc analysis, are indicated with *** showing P ≤ 0.001 and ** showing P ≤ 0.01. (D) Endogenous levels of ABA were determined in after-ripened seeds; *pm19l1* seeds contain about 4 times the level of ABA compared to the wildype. T-test analysis shows a significant difference (**) at P ≤ 0.01 (E) Q- PCR analysis of wildtype (COL), *pm19l1*, *mkk2* and *mkk3* seeds for the transcription factor genes *ABI3*, *ABI4* and *ABI5*. Both *mkk2* and *mkk3* are like *pm19l1*, with reduced *ABI3* and enhanced *ABI4* and *ABI5* levels. (F) Q-PCR analysis of wildtype, *pm19l1*, *mkk2* and *mkk3* seeds for the ABA synthesis gene *NCED6*, the GA synthesis gene *GA20ox1* and the ABA breakdown gene *CYP707A1*. For Q-PCR, Bars show standard deviation. statistically significant differences relative to the wildtype, using Dunnett’s test at a = 0.05, are indicated with an asterisk.

PM19L1 has previously described as a negative regulator of dormancy in Arabidopsis, with loss of function mutants showing higher dormancy in freshly harvested seeds and for heat stress-induced secondary dormancy (Barrero et al., 2019). To explore this further, green siliques at around 3 weeks post fertilisation were harvested, surface sterilised and placed on agar plates at 20 ^0^C. Wildtype seeds germinated after 7 days, whereas germination was much reduced for the immature *pm19l1* seeds, indicative of enhanced dormancy in developing seeds of *pm19l1* (Fig. 7A). To study dormancy in mature seeds, Columbia wildtype and *pm19l1* plants were grown at 20 ^0^C or at 14 ^0^C to induce primary dormancy (Kendall et al., 2011), and freshly harvested seeds put to germinate at 20 ^0^C. There was little difference in germination between wildtype and the mutant when plants were grown at 20 ^0^C, however when grown and matured at 14 ^0^C, the *pm19l1* seeds showed greater dormancy than the wildtype. Both wildtype and mutant responded to 2 days stratification at 4 ^0^C with enhanced germination at 20 ^0^C (Fig. 7B). Dormancy and germination in seeds is associated with the plant hormone ABA, which is responsible for regulating seed development, dormancy, and germination, and for the expression of ABA-induced genes such as the transcription factors *ABI3*, *ABI4* and *ABI5* (Finkelstein, 1994). *ABI3* was downregulated in *pm19-l1* and in dormant seeds of the wildtype, whereas *ABI4* and *ABI5* were found to be upregulated 5-10 fold in *pm19l1* and dormant wildtype seeds (Fig. 7C). However, as *pm19l1* seeds show constitutive upregulation of the dormancy-inducing transcription factor genes *ABI4* and *ABI5*, this cannot be the primary reason for the enhanced dormancy of pm19l1 seeds grown at 14 ^0^C, and another, as yet unidentified, factor must be involved.

**Figure 7.**
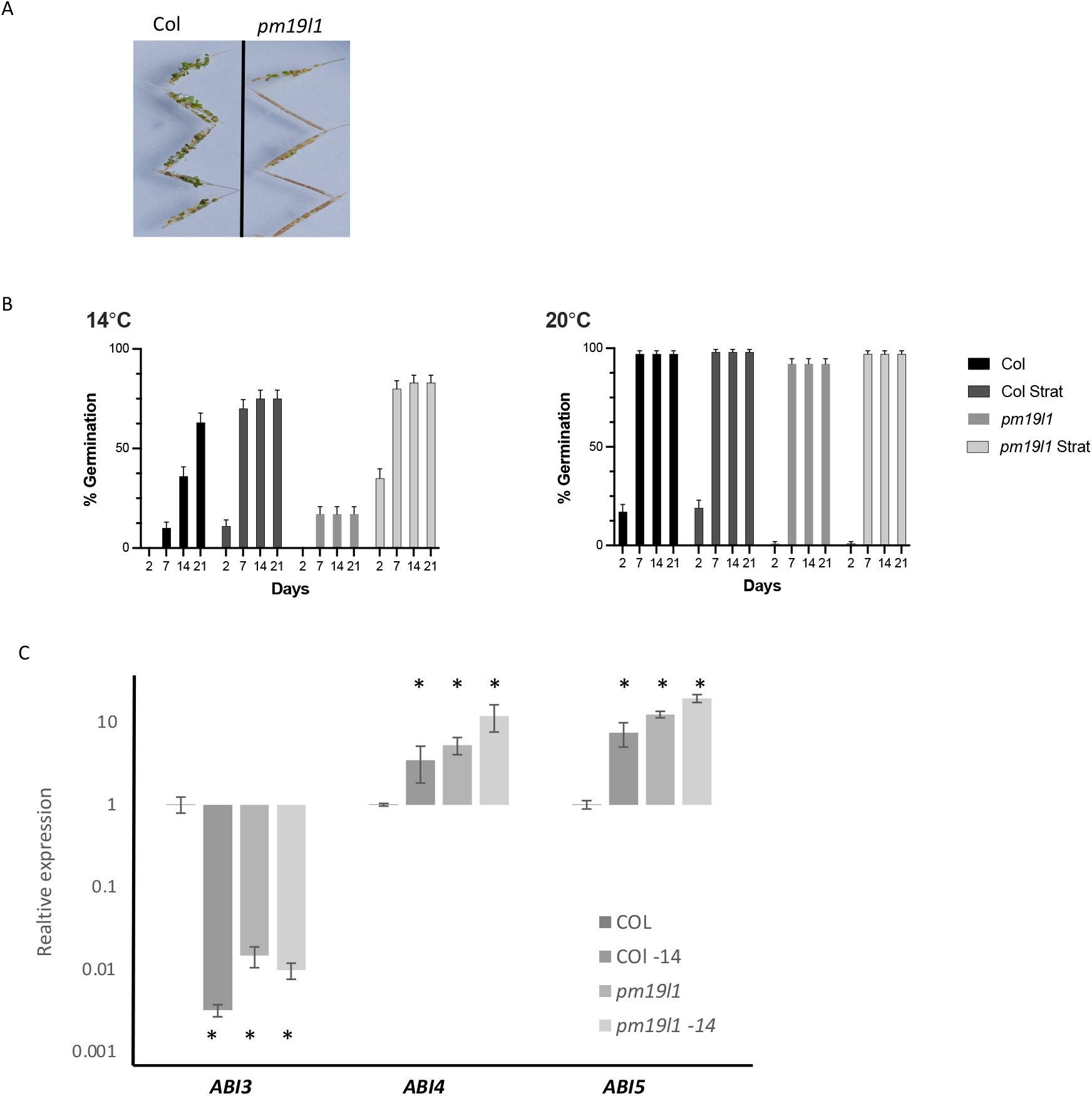
Dormancy analysis of seeds. (A) Green siliques of wildtype Columbia and *pm19l1* at around 2 weeks post fertilisation were incubated on agar plates for 7 days, the *pm19l1* mutant has enhanced dormancy compared to the wildtype. (B) Wildtype Columbia and *pm19l1* plants were grown at 14^0^C to induce dormancy in mature seeds, or at 20 ^0^C. Freshly harvested seeds were then placed on agar plates at 20 ^0^C with or without stratification at 4 ^0^C for 2 days. Bars show standard deviation (based on binomial distribution, 100 seeds). Seeds of *pm19l1* was more dormant than the wildtype control when grown at 14 ^0^C. In all cases, germination was enhanced by stratification. (C) Q-PCR analysis of wildtype (COL) and *pm19l1* seeds grown at 20 ^0^C or 14 ^0^C for transcription factor genes *ABI3*, *ABI4* and *ABI5*. Bars show standard deviation. Statistically significant differences relative to the wildtype, using Dunnett’s test at a = 0.05, are indicated with an asterisk.

## Discussion

The work presented here indicates that the conserved Arabidopsis PM19L1 is an osmosensor that regulates seed dormancy, and also germination under osmotic stress. PM19L1 complements the yeast *sho1* osmosensor mutant. Further, loss of function of PM19L1 results in enhanced dormancy, both in developing and mature seeds, and in enhanced germination sensitivity to salt and osmotic stress. Complementation of yeast osmosensor mutants has been previously used to identify other potential plant osmosensors. The *sln1* mutant (defining the other branch of the Hog1 pathway) is complemented by the Arabidopsis histidine kinase AHK1 (Urao et al., 1999; Tran et al., 2007), and *ahk1* mutants are sensitive to osmolytes during germination (Wohlbach et al., 2008). However, the role of AHK1 in the mature plant has been contested as it is additionally associated with regulation of stomatal density (Kumar et al., 2013). This raises the possibility that PM19L1 and AHK1 fulfil in plant seeds the roles of the two parts of the classic Hog1 osmosensing pathway found in yeast. This parallel extends to the interaction of plant MKK proteins with PM19L1 as demonstrated by both split ubiquitin assays in yeast and, for MKK2 and MKK3, pulldown assays with recombinant proteins and plant protein extracts. Knockout lines of *mkk2* and *mkk3* also showed reduced germination on salt or sorbitol containing media, which suggests that these signalling proteins are part of an osmotic stress signalling pathway together with PM19L1.

The plant hormone ABA is implicated in the control of seed dormancy and germination, and analysis of the *pm19l1* mutant shows that PM19L1 acts through modulating ABA signalling. Levels of *ABI4* and *ABI5* gene expression are enhanced in mature *pm19l1* seeds, indicating that these transcription factor genes are negatively regulated through PM19L1. This suggests a possible mechanism for the deeper dormancy and salt sensitivity seen for *pm19l1*. ABI4 controls seed development and germination by enhancing sensitivity to ABA (Finkelstein et al., 2002) and as a positive regulator of ABA biosynthetic genes and a negative regulator of genes involved in ABA inactivation and GA biosynthesis (Shu et al., 2013, 2016). This is consistent with our findings that for *pm19l1*, endogenous ABA levels are enhanced, *NCED6* expression is upregulated, and *CYP707A1* expression downregulated, that paclobutrazol has a greater impact on germination of *pm19l1* than of the wildtype, expression of *GA20ox1* is reduced in *pm19l1*, and that exogenous GA will enhance germination of *pm19l1* on salt. A further mechanism by which ABI4 may act as an attenuator of germination is because ABI4 promotes ROS production through the expression of RbohD and hence represses germination (Luo et al., 2021). *ABI5* expression is promoted by ABA and also by ABI4; ABI5 enhances sensitivity to ABA, leading to inhibition of germination and seedling growth arrest, whereas *abi5* mutants are salt insensitive (Lopez-Molina et al., 2001; Skubacz et al., 2016). Since both *ABI4* and *ABI5* gene expression was enhanced in the *mkk2* and *mkk3* mutants, this suggests a pathway in which the osmotic environment is perceived through PM19L1 and the signal transduced through MKK2 and MKK3 to modulate *ABI4* and *ABI5* expression, resulting in a reduction of ABA and an increase in GA levels in the seed.

In contrast to *ABI4* and *ABI5*, levels of *ABI3* were markedly reduced in mature *pm19l1* seeds. As well as contributing to the ABA signalling pathway, ABI3 (together with LEC1, LEC2 and FUS3), is a transcription factor that regulates the transition between early and late embryo and seed development and germination (Parcy et al., 1997; Jia et al., 2014). Severe *abi3* mutants are characterised by a failure to transition to the mature phase of seed development, remaining green and desiccation intolerant, and with reduced dormancy (Nambara et al., 1995), and although this extreme phenotype was not observed in *pm19l1* since some level of ABI3 transcript was still detectable, it is possible that the transition mediated by ABI3 is in part triggered by the osmotic environment of the desiccating seed through PM19L1. The same molecular phenotypes, upregulation of *ABI4* and *ABI5* expression, and downregulation of *ABI3* expression was seen for seeds of *mkk2* and *mkk3* mutants, which further strengthens the argument for an osmoregulatory pathway consisting of PM19L1 and MKK proteins.

The wheat PM19L1 homologues *PM19-A1* and *PM19-A2* have been identified as QTL candidate genes for preharvest sprouting and dormancy (locus Phs1 on chromosome 4A, (Barrero et al., 2015)), with expression of these genes being higher in dormant accessions. Additionally, other dormancy QTL genes in both wheat and barley have been identified as being homologues of *MKK3*, the wheat *MKK3*, also mapping to QTL Phs1 (Nakamura et al., 2016; Torada et al., 2016). This indicates that in cereals, PM19-A1/PM19-A2 and MKK3 all contribute to dormancy and preharvest sprouting, consistent with a signalling pathway that leads from the osmosensor through a MAP kinase cascade. The rice PM19L1 homologue OSPM1, which is present as multimeric forms in the membrane, has previously been found to facilitate ABA influx into yeast (Yao et al., 2018), which may also be potential mechanism that facilitates control of germination and dormancy. Lastly, the evolutionary ancient roots of PM19L1 may represent an important adaptation to terrestrial life for plants in regulating the plant response to osmotic stress.

## Materials and Methods

### Plant material and growth conditions

The Arabidopsis plants used in this study were the wildtype (accession Columbia, Col-0), the homozygous T-DNA insertion mutants *pm19l1* (line SALK_075435) in the Col-0 background, and transgenic lines bearing the constructs *pCambia1302, pCaMV35S:PM19L1-GFP* and *pPM19L1:PM19L1* in both wildtype and *pm19l1*mutant backgrounds, and *pPM19L1:UidA* in the wildtype background. In addition, T-DNA insertion mutants *mkk1,* (Salk line 027645) (Qiu *et al*., 2008), *mkk2* (SAIL_511H01) (Qiu *et al*., 2008) and *mkk3* (SALK_051970) (Takahashi *et al*., 2007) were studied. Seeds were harvested from single plants of homozygous lines pot-grown in a mixture of peat (80%) and vermiculite (20%) in a growth chamber with the above-mentioned conditions. Unless otherwise indicated, seeds were after-ripened at room temperature for at least 2 weeks.

### Phylogenetic tree construction

An existing alignment from Yao *et al*., (2018) was used and additional sequences added. The Arabidopsis PM19L1 protein sequence was used for BLAST searches on proteomes available on Phytozome and other sources (see Tables S1, 2). No E value cut off was used and all possible sequences were investigated. New protein sequences were added to the existing alignment using mafft-add (MAFFT version 7). The alignment was trimmed to include only regions of the predicted transmembrane domains as illustrated in Fig. 1B. This trimmed alignment was then used to construct the phylogenetic tree using PhyML 3.0. The LG protein substitution model, nearest neighbour interchange for tree searching and approximate likelihood ratio (SH-like) branch support settings were used.

### Binary plasmid construction and plant transformation

To generate the *pCaMV35S:PM19L1-GFP* lines, the full-length ORF of *PM19L1* (AT1G04560) from cDNA clone *PAP111* (Genbank Z29867) was subjected to site-directed mutagenesis using primers PM19-SDM1 and PM19-SDM2 (Table S5) to remove an *Nco*I site from the coding region, and the coding region amplified with primers PM19-NcoI (F) and PM19-Spe1 (R) so as to introduce an *Nco*I site at the encoded N-terminal and a *Spe*I site at the C terminal. This amplicon was cloned into the pCambia1302 vector over *Nco*I and *Spe*I and checked by sequencing to yield the *pCaMV35S:PM19L1-GFP* construct. The construct was transferred into *Agrobacterium tumefaciens* strain EHA105, as was pCambia 1302 as a control, and used to produce transgenic plants by the floral dip method (Clough and Bent, 1998). Transgenic plants were screened for hygromycin resistance, and homozygous T3 generation seeds were selected.

To generate the *pPM19L1:PM19L1* complementation lines, a 2894 bp fragment was amplified from wildtype Arabidopsis genomic DNA using primers PM19-3 (F) and PM19-4A (R) using Phusion proofreading enzyme. The fragment was digested with *Eco*RI and ligated into binary vector pBinHyg (Qiu *et al*., 2008), digested with *Eco*RI and *Sma*I, which replaces the CaMV35S promoter in the vector. This clone contained 1970 bp of sequence upstream to the *PM19L1* translational start site, the *PM19L1* gene including introns, and 165 bp of genomic sequence upstream to the *PM19L1* translational stop site. Transgenic plants were screened for hygromycin resistance, and homozygous T3 generation seeds were selected. To generate the *pPM19L1:UidA* construct, a 2282 bp fragment was amplified from wildtype Arabidopsis genomic DNA using primers PM19-3 (F) and PM19-SDM2(R) and Phusion proofreading enzyme, and cloned into pGEM T-Easy. This clone contained 1970 bp of sequence upstream to the *PM19L1* translational start site, and a further 312 bp of coding sequence. A 1257 bp *Eco*RI-*Bgl*II genomic fragment (*Bgl*II is in the 5’ untranslated region of *PM19L1*) was excised and cloned into the binary vector pPR97 in front of the *UidA* gene (Szabados *et al*., 1995). Transgenic plants were screened for kanamycin resistance, and homozygous T3 generation seeds were selected.

### Germination assays

Sterilized seeds were sown on 0.8% agar media with half-strength MS (0.5× MS salts, 10 mM MES pH 5.6) and additives as indicated in Figs. 3 and 6. Plates were placed in a growth chamber with 8/16 h light/dark cycle, with 200 µE m-2 s-1 white light, 22/20 °C light/dark temperature, and 70% relative humidity. Germination was scored for by observing radicle protrusion. 100 seeds were used per assay and analysed using binomial statistics. For stress treatments, seedlings were grown for 14 days on half-strength MS medium plates. Seedlings were then sprayed directly with either a 15% w/v solution of PEG 6000, 25 µM ABA, 200 mM NaCl, 200 mM KCl, or distilled water and incubated for 24hrs. They were frozen in liquid nitrogen in pools of 5 seedlings per line in triplicate, ready for RNA extraction and mRNA expression. For precocious germination assays, developing siliques at the long-green stage were collected, slightly opened at the replum-valve margin using a surgical blade, sterilized with 70% (vol/vol) ethanol for 1 min and 25% bleach for 10 min, and then plated on half-strength MS medium plates.

### Physiological measurements

Measurement of free proline content was carried out according to Bates *et al*., (1973). Soluble sugars were measured according to Yemm and Willis, (1954) using an anthrone reagent. In both cases, triplicate extracts were prepared and measured. Data was analysed for statistical significance by two-sided ANOVA and post-hoc Tukey’s test for multiple comparisons. Potassium and sodium ion measurements were carried out on aqueous plant extracts using Laqua twin ion-specific electrodes (Horiba Scientific). ABA was quantified by HPLC using an Athena 4.6 x 250 mm 5 µm C18 column as described by Nakurte *et al*., (2012), with C18 SPE based extraction of triplicate 0.75 gram seed samples and with cinnamic acid added as an internal standard.

### Histochemical staining for B-glucuronidase (GUS) assay and GFP imaging

Histochemical GUS staining was carried out on seeds and seedlings of pPM19L1:UidA lines, grown on agar plate media as described by Jefferson, (1987). After incubation, the pigments were removed by repeated washing in 70% (v/v) ethanol. Images of GUS-stained seedlings were recorded with a Leica DFC320 microscope. Imaging of GFP-expressing stable Arabidopsis lines was performed using an inverted ZEISS LSM880 equipped with a C-Apochromat 40x/1.2 W Korr W27 water immersion objective. Prior to imaging, the seedlings were incubated in 10 µM FM4-64 for 10 min and mounted in water. GFP as well as FM4-64 were excited at 488 nm and GFP was detected at 510-550 nm while FM4-64 was detected at 660 - 740 nm. To avoid cross talk, the two fluorophores were acquired separately.

### RNA extraction and mRNA expression

Total RNA extraction and cDNA synthesis were performed as described by Alexander *et al*., (2019). Northern blotting was carried out at described by Alzwiy and Morris, (2007). Plant material was taken from agar-grown seedlings or mature, soil-grown plants as described above. Dormant and active axillary buds were prepared by excising nodal stem segments and placing into microcentrifuge tubes containing liquid half MS for 3 days using a modified protocol described in Seale *et al*., (2017). Dormant buds were harvested from intact stem segments and active buds harvested after removing the primary shoot apex to activate buds 24 hours previously. Oligonucleotides were designed using Primer3plus (Untergasser *et al*., 2012) (Table S5). RT-qPCR was performed with a StepOnePlus™ Real-Time PCR System (Applied Biosystems). Standard cycling conditions were 10 min at 95 °C followed by 40 cycles of 15 s at 95 °C and 1 min at 60 °C; the melting curve profiles were then determined. Reactions were performed in three technical replicas per biological replica and in 3 biological replicas per line and treatment. Expression values were normalized to the Arabidopsis *Actin-2* (*ACT2*, At3g18780) and gene expression values were calculated using the 2^−ΔΔ*C*T^ method (Schmittgen and Livak, 2008).

### Yeast strains and plasmids

All strains and plasmids used for work in the yeast *Saccharomyces cerevisiae are* described in Supplementary Tables S3 and S4. The ORF from *PM19L1* cDNA clone *PAP111* (Cooke *et al*., 1996) was amplified with primers PM19-3A and PM19-Xba1 to yield a full-length ORF which was cloned into the *Eco*RV site of pBluescript. *PM19L1* was re-excised with *Bam*HI and *Xba*I and cloned into pYES2 to yield pPMRA1, which was used to test for complementation of *trk1/trk2* (Bertl *et al*., 2003) and of *ena1* (Haro *et al*., 1991).

To test for complementation of the YSH642 strain (*gdp1/gdp2*) (Pettersson *et al*., 2006), auxotrophic selection over leucine was required, therefore the LEU2 expression cassette from pGAD424 was cloned into the *Apa*I and *Nhe*I sites in the URA3 gene of pYES2 to give pPMRA2, and into pYES2-PM19 to give pPMRA3.

In order to test for complementation of the *sho1* mutant (Maeda *et al*., 1995), auxotrophic selection over histidine was required, therefore the HIS3 expression cassette from pFA6a- HIS3MX6 was cloned into the *Apa*I and *Nhe*I sites in the URA3 gene of pYES2 to give pPMRA4, and into pYES2-PM19 to give pPMRA5. A shortened version of *PM19L1*, which removed the non-conserved C-terminal was created by use of primers PM19-HindIII and Short PM19- XbaI to yield an amplicon encoding a protein of 149 amino acids (PM19L1**Δ**) that was cloned into pPMRA4 to give pPMRA6.

To additionally express Arabidopsis *MKK* genes together with *PM19L1* in the *sho1* mutant, selection over kanamycin was employed, therefore the KAN expression cassette from pFA6a-KANMX6 was cloned into the *Apa*I and *Nhe*I sites in the URA3 gene of pYES2 to give pPMRA7. Subsequently the open reading frames of *AtMKK1* (At4g26070), *AtMKK2* (At4g29810) and *AtMKK3* (At5g40440) were cloned into pPMRA7 to give pPMRA8, pPMRA9 and pPMRA10 respectively. Site directed mutagenesis was used to change specific amino acids in PM19L1 and these modified versions were introduced into pPMRA4 to give pPMRA11 (79W-G), pPMRA12 (80R-G), and pPMRA13 (82D-G). The Arabidopsis tetraspanin 3 (At3g45600) ORF was amplified using primers TET3-F and TET3-R and cloned into pPMRA4 to give pPMRA14.

### Yeast growth conditions

Yeast cells were grown using standard microbial techniques and media (Amberg *et al*., 2005). Media designations are as follows: YPAD is 1% Yeast Extract - 2% Peptone - 2% glucose plus Adenine medium; SDAP is synthetic dextrose arginine phosphate (Rodríguez-Navarro and Ramos, 1984). For YP-Gal, glucose was replaced with 2% galactose. SD is Synthetic Defined dropout (SD-drop-out) medium. Minimal dropout media are designated by the constituent that is omitted (e.g. -leu –trp –his –ade -ura medium media lacks leucine, tryptophan, histidine, adenine and uracil). Recombinant plasmid DNA constructs were introduced using the standard lithium acetate method (Gietz and Schiestl, 2007). Selection and growth of transformants was performed in SD -drop-out lacking the appropriate amino acid or containing the appropriate antibiotic.

### Functional complementation of yeast mutants

For complementation analysis all transformed yeast cells were grown in selective media and collected at a density of OD 0.4-0.6. Subsequent fivefold dilutions were made and 5 μl were spotted onto minimal medium plates containing galactose as a carbon source and the indicated osmotica. They were incubated at 30°C (unless stated) for 2–4 days and scanned. Potassium and sodium-dependent phenotypes were analysed using the PLY240 and L5709 strains (Table S3) as described in Bertl *et al*., (2003) and Haro *et al*., (1991) respectively. For potassium importer assays, wildtype (JRY379) and *trk1/2* (PLY240) strains transformed with pYES2 or pPMRA1 were grown on SDAP agar plates (pH 5.8), supplemented with 20 mM KCl. For sodium assays wildtype (W303-1A) and *ena1* (L5709) strains were transformed with pYES2 or pPMRA1 and grown on SD-ura plates supplemented with of NaCl at 500 mM. Aquaglyceroporin function was tested by transforming the wildtype (W303-1A) strain or YSH642 (Pettersson *et al*., 2006) (Table S3) with pPMRA2 or pPMRA3 and growing them on SD-L plates supplemented with KCl or sorbitol. The *sho1*Δ, *ssk2*Δ, *ssk22Δ* triple mutant (TM310) (Maeda *et al*., 1995) was transformed with the pPMRA4 control vector, pPMRA5 and pPMRA6, or mutant variants pPMRA11-13 (Table S4). Osmosensitivity of yeast strains was tested by growth on SD-H (Gal) plates with and without different concentrations of NaCl or sorbitol. The *sho1*Δ, *ssk2*Δ, ssk22Δ triple mutant containing pPMRA5 or pPMRA6 was then re-transformed with Arabidopsis *MKK1*, *MKK2* or *MKK3* in the yeast vector pPMRA7 containing a kanamycin resistance marker (plasmids pPMRA8-10). These strains were tested by growth on SD-H (Gal) / G418 plates, with and without different concentrations of sorbitol as described in the figure legend (Fig. 4).

### Yeast two hybrid screens

Protein-protein interaction analysis was carried out by utilising the split-ubiquitin yeast two-hybrid system (Snider *et al*., 2010). All work was done using strain NMY51 (Dualsystem Biotech, Table S3). NMY51 was first transformed with the pAMBV bait plasmid containing one of 6 different PM19L1 variants (pPMRA15 - 20, Table S4). Auto-activation was determined by co-expressing each bait with NubI ‘positive’ and NubG ‘negative’ control prey plasmids. Ideally, all transformed strains would be expected to show comparable growth on the transformation selection medium, but only bait strains containing NubI ‘positive’ control prey would be expected to grow on the interaction selection medium. Baits containing NubG ‘negative’ control preys that grow on interaction selection medium are said to be ‘self-activating’ and are not suitable for screening. All constructs proved suitable for screening. Prey vector pPR3C containing one of the full-length *MKK1*, *MKK2* and *MKK3* sequences (pPMRA21-23, Table S4) were then co-transformed with strains containing the bait plasmids. Transformants were selected for the presence of both bait and prey plasmids during 3 days of growth at 30°C on SD-trp-leu medium. Positive colonies were transferred to liquid SD-trp-leu medium and grown overnight to an OD of 1.0. Five microliters of different dilutions (1:10, 1:100, 1:1000) were spotted onto transformation selection medium (SD-trp-leu, to ensure that spotted cells are carrying the appropriate plasmids) and onto interaction selection medium (SD-trp-leu-his-ade media and SD-trp-leu -Xgal, to select specifically for cells containing interacting bait–prey pairs). Plates were then incubated for 4 -6 days at 30°C.

### Pulldown assays

Recombinant proteins were produced as maltose binding protein fusions. MBP-MKK1 has been described in (Huang *et al*., 2000). MBP-MKK2 and MBP-MKK3 were produced in a similar fashion by PCR amplifying and cloning the open reading frame of each gene (primers MKK2-EcoRI_F, MKK2-SalI_R, MKK3-XbaI_F, MKK3-PstI_R) into the pMAL-C2 vector and fusion proteins were expressed and purified using amylose-affinity chromatography as described by the manufacturer (New England Biolabs). The over-expressed recombinant proteins were then coupled to CNBr-activated Sepharose (Sigma) at a ratio of 2.5 mg protein to 1 ml Sepharose slurry as described by the manufacturers.

Crude protein extracts from 6-day old transgenic lines bearing the constructs *pCaMV35S:PM19L1-*GFP and *pCaMV35S:* GFP were prepared as described in Kadota *et al*., (2016). Briefly plant material was ground in liquid nitrogen and added to extraction buffer (50 mM Tris-HCl pH7.5, 150 mM NaCl, 10% glycerol, 1 % IGEPAL, 100 mM Na_2_MoO_4_, 1 mM NaF, 1.5 mM Na_3_VO_4_, 25 mM PMSF). For pull-down assays, equal amounts of protein extracts were incubated with 100 µl MKK column matrices for 1 h at room temperature followed by five washes with 1 ml wash buffer (50 mM Tris pH 7.5, 50 mM NaF, 500 mM NaCl). Proteins were eluted from the matrix by resuspension in denaturing SDS-PAGE sample buffer. Total soluble proteins were then separated by 10% SDS-PAGE and transferred to a nitrocellulose membrane. Blotted membranes were blocked with PBS containing 4% w/v skimmed milk powder followed by incubation with 1:1000 dilution of mouse anti-GFP antibody (GF28R, Invitrogen) in blocking buffer followed by incubation with 1:5000 dilution of goat anti-mouse IgG secondary antibody, HRP (A4416, Sigma-Aldrich) and detected by chemiluminescence.

## Author Contributions

Ross Alexander carried out yeast complementation, yeast split ubiquitin assays and Q-PCR work, and wrote the manuscript. Pablo Castillejo-Pons carried out mutant phenotype analysis and produced plant transformation constructs, Omar Alsaif carried out mutant phenotype analysis and plant transformations, Vivien Strotmann carried out GFP imaging, Yvonne Stahl carried out gene expression studies and helped with protein-protein interaction work, Madeleine Seale carried out gene expression studies and protein phylogeny, Peter Morris conceived the project, designed experiments, carried out plant transformations and mutant phenotype analysis and wrote the manuscript.

## Data and materials availability

All data is available in the main text or the supplementary materials.

---

**Figure S1.**
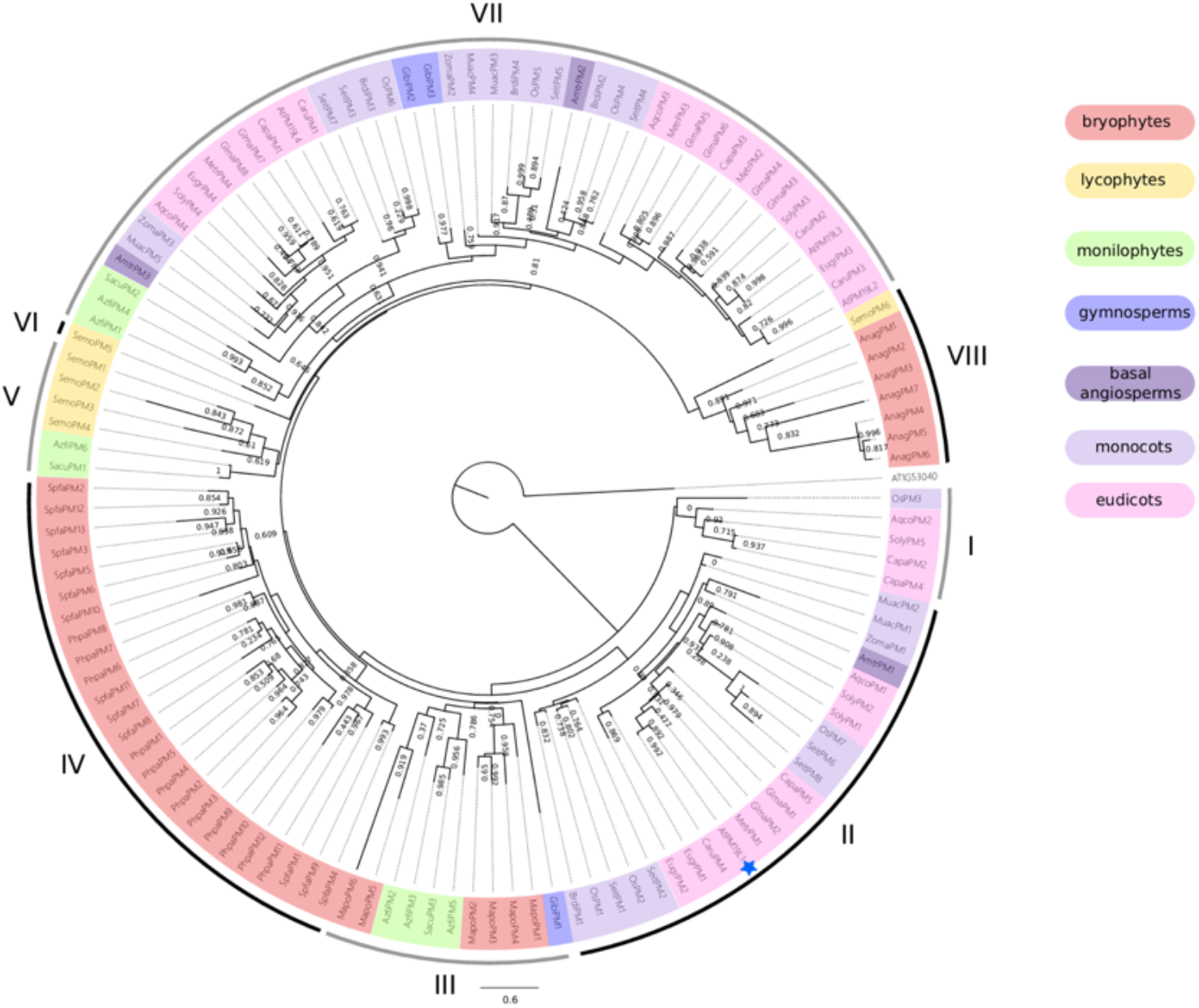
Phylogenetic tree indicating the relationship between AWPM-19 family members across land plants. Scale bar indicates mean number of amino acid substitutions per site. Node numbers indicate statistical support for that node configuration according to the approximate likelihood ratio test (aLRT) method. The protein sequence alignment from ^7^ was used with additional species added here. An Arabidopsis protein, AT1G53040 exhibiting weak sequence similarity to AWPM-19 proteins was used as an outgroup to root the tree. Coloured shading illustrates taxonomic divisions and roman numerals indicate monophyletic clades. AtPM19L1 is indicated with the blue star. See Table S1 for details of species and gene IDs. The phylogeny demonstrates that the AWPM-19 gene family evolved when land plants originated (no similar sequences were found in any algae). Class VII AWPM-19 genes appear to be conserved from ferns to angiosperms, while class II, including *AtPM19L1* is an angiosperm specific clade. There are also many AWPM-19 in bryophytes and some in lycophytes and ferns that have diverged from one another into separate clades and are not generally conserved among seed plants.

**Figure S2.**
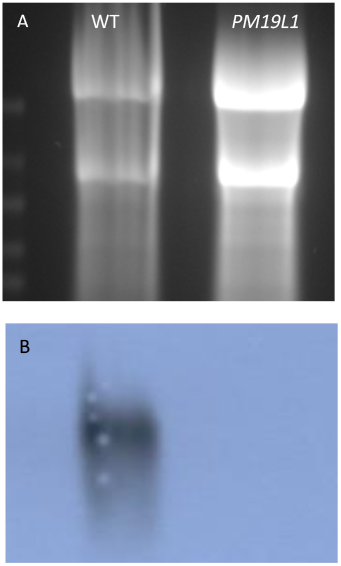
Northern blot analysis of total RNA isolated from wildtype and *pm19l1* seeds and hybridised with a digoxygenin labelled *PM19L1* probe. (A) Ethidium-stained RNA, (B) chemiluminescence signal from hybridised probe. There is no detectable signal for *PM19L1* in the mutant.

**Figure S3.**
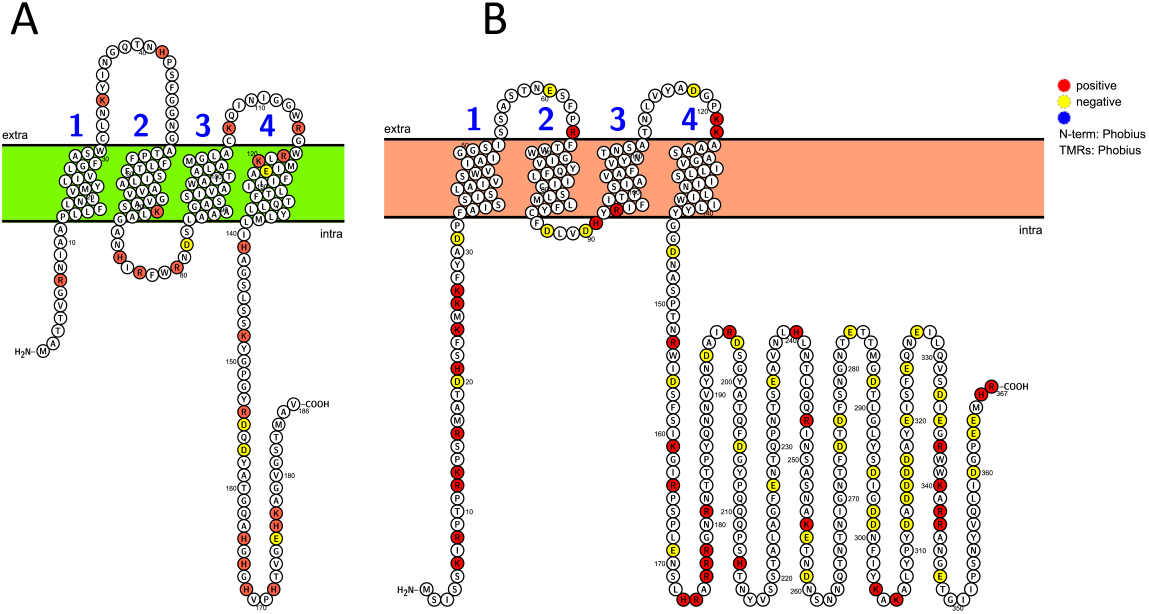
Similarity in predicted secondary structure for (A) PM19L1 and (B) Sho1, carried out using Protter (http://wlab.ethz.ch/protter/start/)

**Figure S4.**
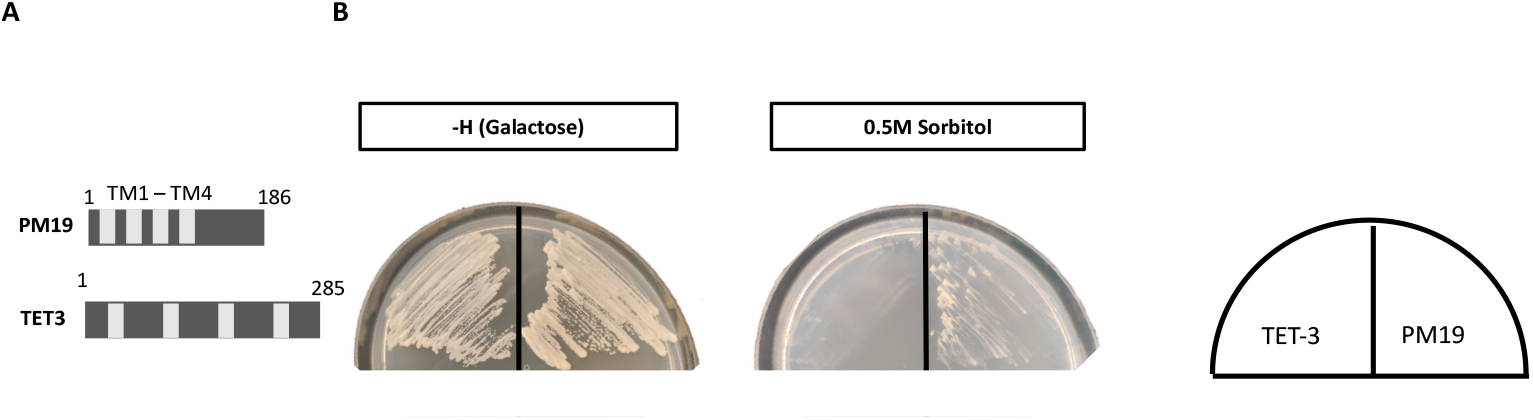
Transformation of yeast osmosensing *ssk1/ssk22/sho1* mutant with Arabidopsis tetraspanin (TET-3) and with *PM19L1 (*PM19), showing lack of phenotypic complementation by tetraspanin on media with high osmotic potential. (A) shows the predicted transmembrane domains in each protein. (B) shows growth of *ssk1/ssk22/sho1* transformed with *TET3* or *PM19L1* on media with and without addition of 0.5 sorbitol.

**Figure S5.**
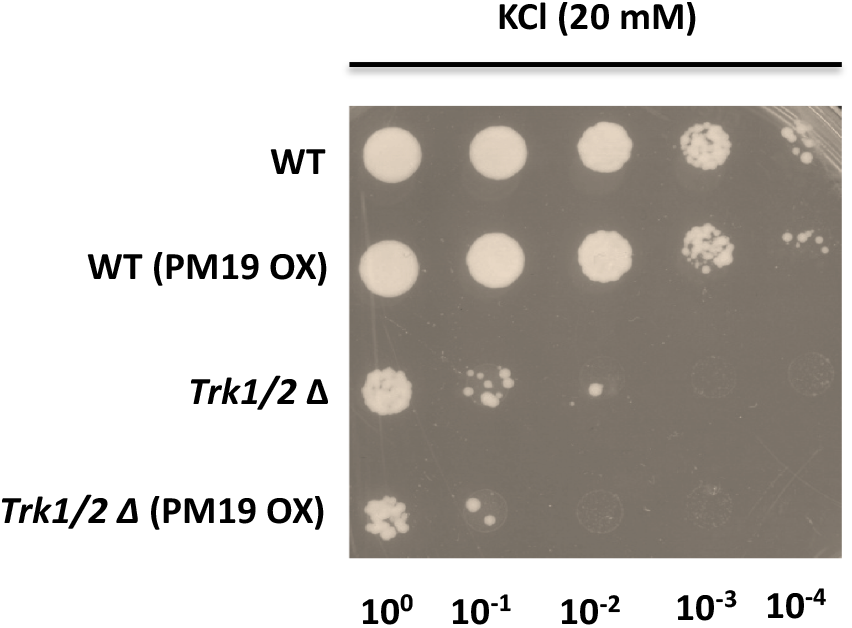
Transformation of yeast potassium transporter mutant *trk1/trk2* with Arabidopsis *PM19L1* in plasmid pPMRA1, showing lack of phenotypic complementation on media with low potassium content. Wildtype strain JRY379 (WT) and *trk1/2* mutant strain PLY240 were transformed with empty vector pYES2 or with pPMRA1 (PM19L1 in pYES2, termed PM19 OX in the figure) and were grown on SDAP agar plates with 20 mM KCl.

**Figure S6.**
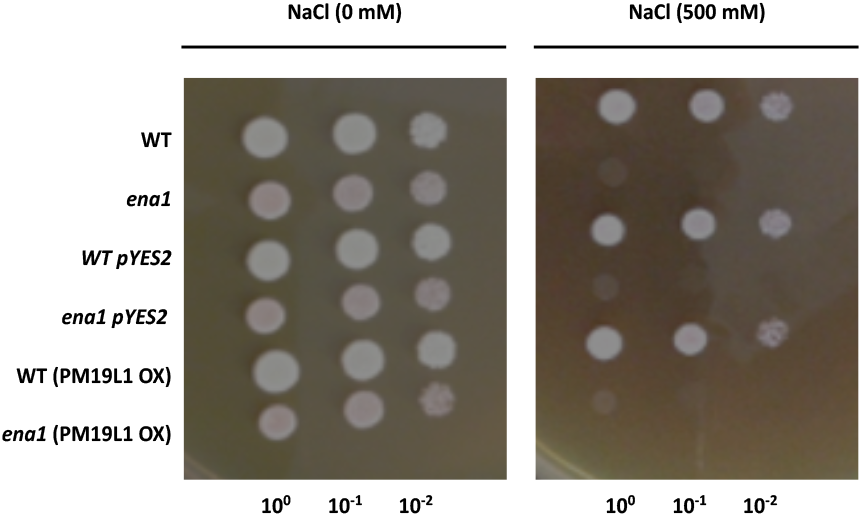
Transformation of yeast sodium transporter mutant *ena1* with Arabidopsis *PM19L1* in plasmid pPMRA1 showing lack of phenotypic complementation on media with high NaCl. Wildtype (WT, W303-1A) and *ena1* (L5709) strains were transformed with pYES2 or pPMRA1 (PM191 OX) and grown on SD-ura plates supplemented with NaCl at 500 mM.

**Figure S7.**
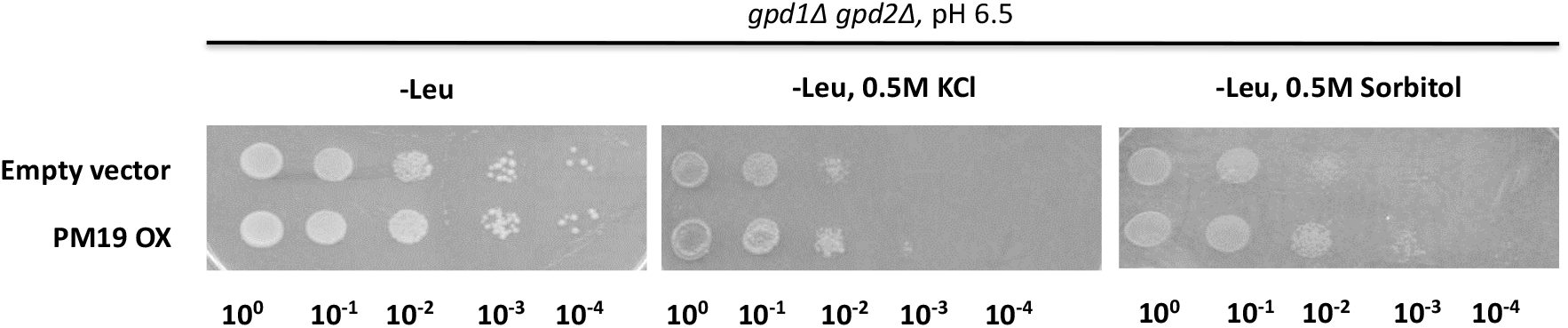
Transformation of yeast glycerol biosynthetic mutant *gpd1/gpd2* with Arabidopsis *PM19L1* in plasmid pPMRA3 (PM19 OX), showing lack of phenotypic complementation when strains were grown on high osmotic strength media with 0.5 Molar KCl or 0.5 Molar sorbitol.

**Figure S8.**
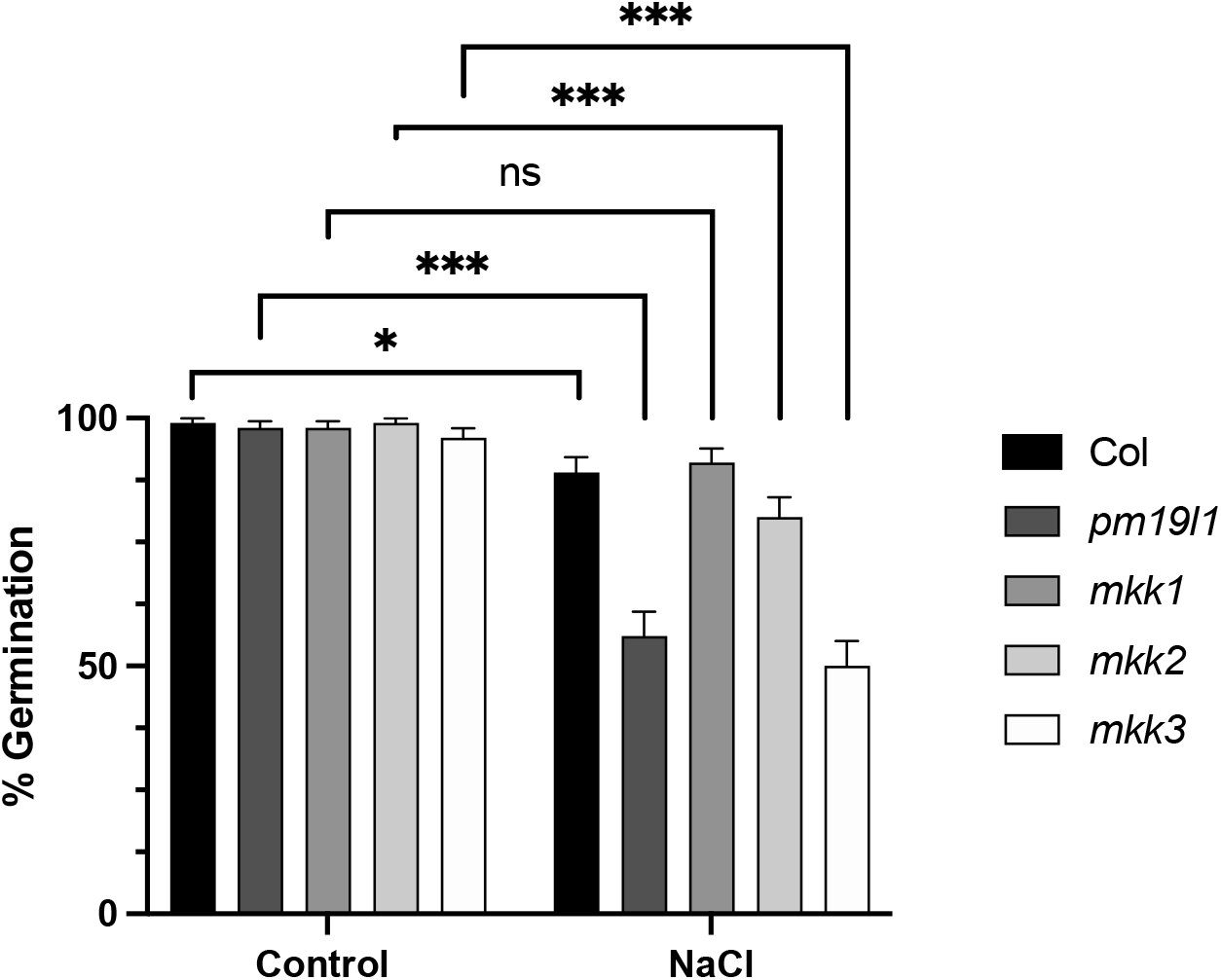
Germination after 7 days of wildtype (Col), *pm19l1*, *mkk1*, *mkk2* and *mkk3* seeds on MS media with and without inclusion of 140 mM NaCl. Bars show standard deviation (based on binomial distribution, 100 seeds). Statistically significant differences, using Fisher’s exact test followed by a Bonferroni-Šídák post-hoc analysis are indicated with ns indicating no significance, * showing P≤ 0.05 and *** showing P ≤ 0.001.

**Figure S9.**
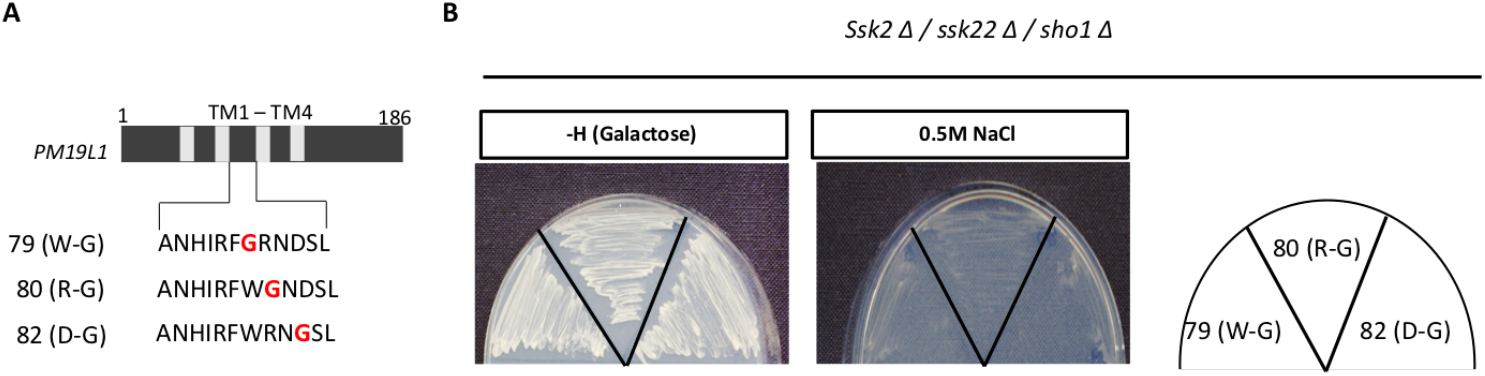
Lack of complementation of *sho1* when conserved amino acids in intracellular domain of PM19L1 are modified. (A) shows the modifications in the intracellular loop between transmembrane domains 2 and 3. (B) shows growth of *skk1/skk2/sho1* transformed with *PM19L1* (pPMRA5) with site directed mutations of specific amino acids as indicated, on galactose-containing growth media without or with addition of 500 mM NaCl to the medium.

**Figure S10.**
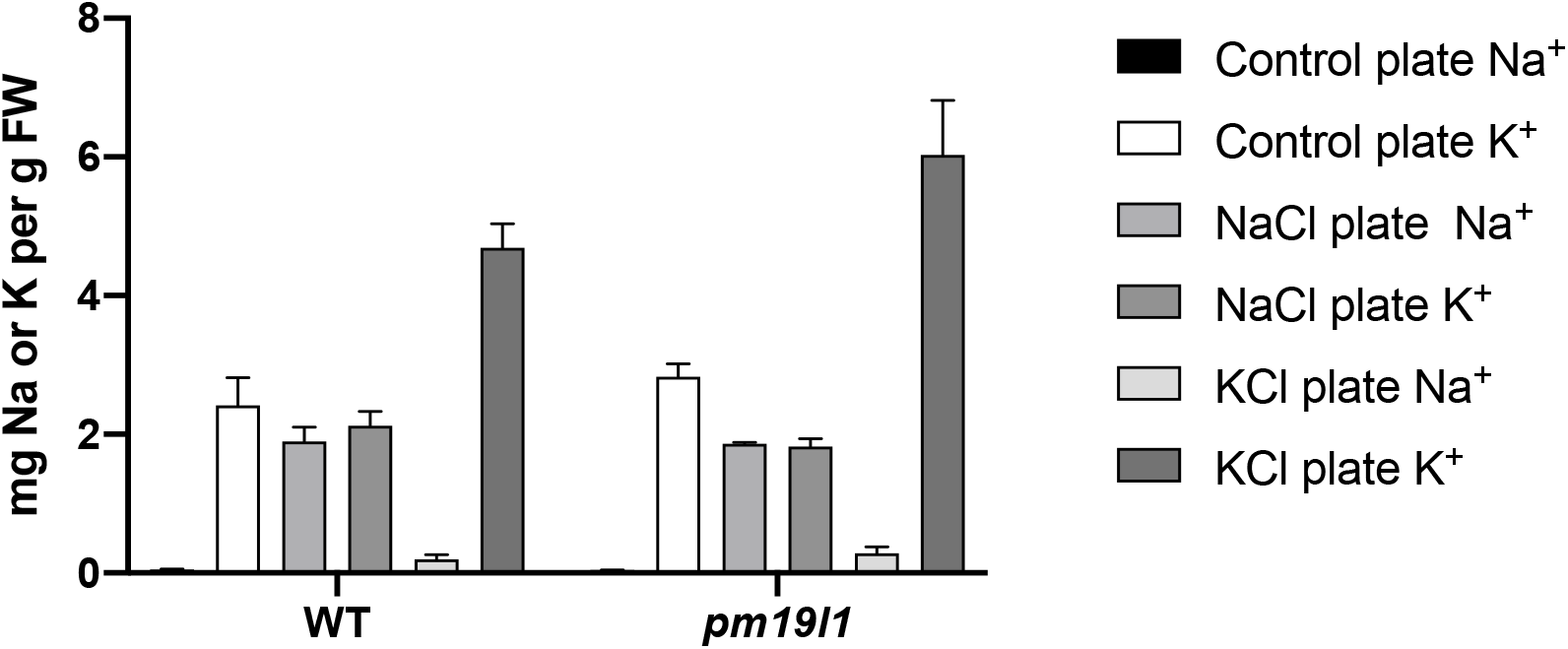
Sodium and potassium ion content of wildtype and *pm19l1* mutant seedlings grown for one week on 60 mM NaCl or KCl. Extracts were prepared from triplicate plates. Bars show standard deviation. ANOVA and post-hoc Tukey’s test for multiple comparisons showed no significant differences between the wildtype and *pm19l1* for all treatments.

**Figure S11.**
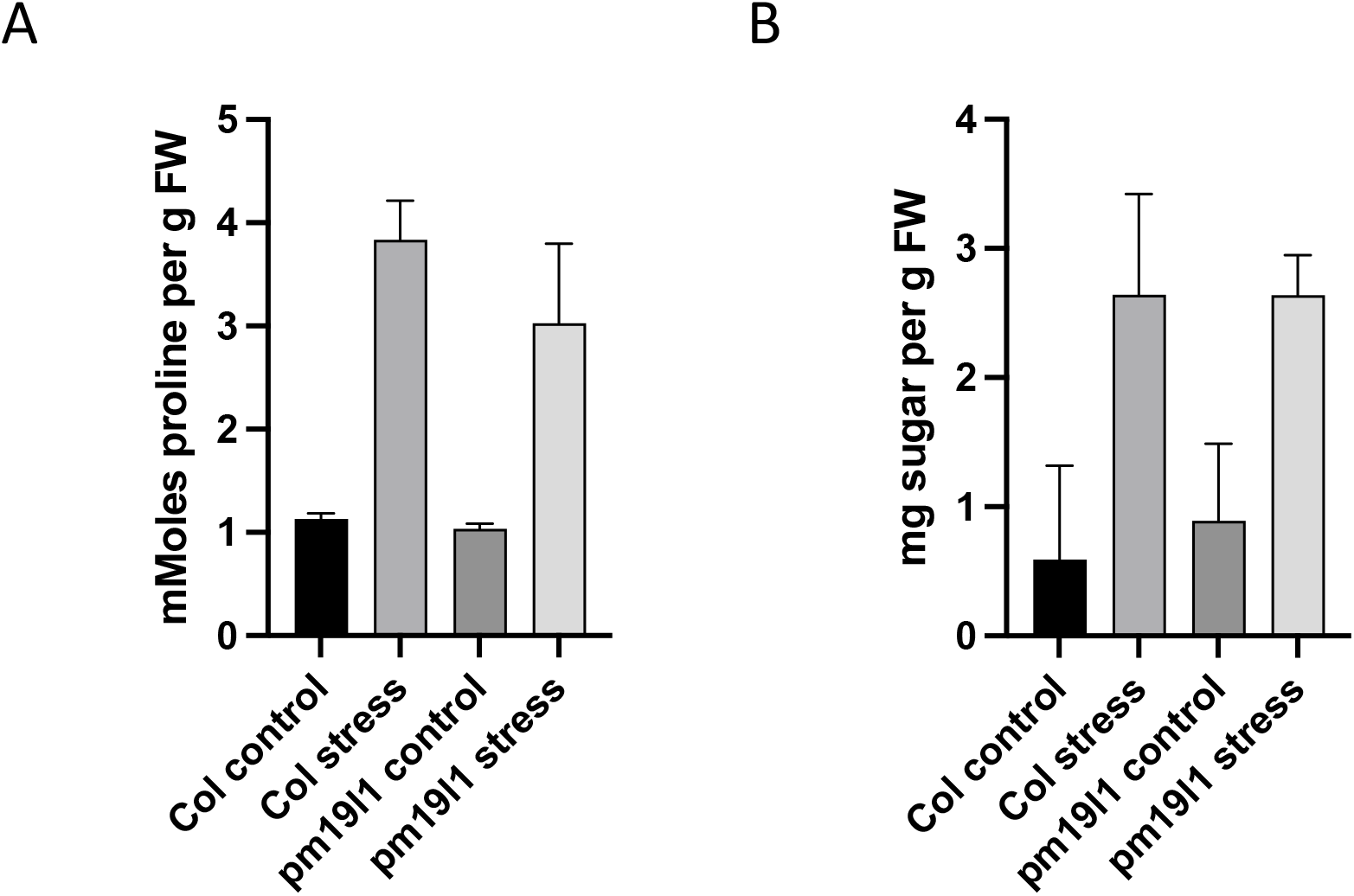
(A) Proline and (B) soluble sugars content of wildtype and *pm19l1* mutant seedlings grown for one week on control or 60 mM NaCl containing media. Extracts were prepared from triplicate plates. Bars show standard deviation. ANOVA and post-hoc Tukey’s test for multiple comparisons showed no significant differences between the wildtype and *pm19l1* for all treatments.

**Table S1.** Number of AWPM-19 encoding genes in different plant species, indicating which were added to the existing alignment from Yao *et al*. (2018) to create the phylogeny seen in Fig. S1.

| Species | Source | Genome/proteome version | No. proteins found |
| --- | --- | --- | --- |
| *Marchantia polymorpha* | Phytozome | v3.1 | 6 |
| *Sphagnum fallax* | Phytozome | v0.5 | 13 |
| *Zostera marina* | Phytozome | v2.2 | 3 |
| *Setaria italica* | Phytozome | v2.2 | 8 |
| *Brachypodium distachyon* | Phytozome | v3.1 | 4 |
| *Solanum lycopersicum* | Phytozome | iTAG2.4 | 5 |
| *Eucalyptus grandis* | Phytozome | v2.0 | 4 |
| *Carica papaya* | Phytozome | ASGPBv0.4 | 5 |
| *Capsella rubella* | Phytozome | v1.0 | 4 |
| *Glycine max* | Phytozome | Wm82.a2.v1 | 8 |
| *Medicago truncatula* | Phytozome | Mt4.0v1 | 4 |
| *Azolla filiculoides* | fernbase | v1.1 | 6 |
| *Salvinia cucullata* | fernbase | v1.2 | 3 |
| *Anthoceros agrestis* | UniZurich | Oxford | 7 |
| *Physcomitrium (Physcomitrella) patens* | Yao *et al.* 2018 |  | 12 |
| *Selaginella moellendorffii* | Yao *et al.* 2018 |  | 6 |
| *Ginkgo biloba* | Yao *et al.* 2018 |  | 3 |
| *Amborella trichopoda* | Yao *et al.* 2018 |  | 3 |
| *Aquilegia coerulea* | Yao *et al.* 2018 |  | 4 |
| *Arabidopsis thaliana* | Yao *et al.* 2018 |  | 4 |
| *Musa acuminata* | Yao *et al.* 2018 |  | 5 |
| *Oryza sativa* | Yao *et al.* 2018 |  | 7 |

**Table S2.** Gene IDs for PM19-encoding genes added to the existing alignment from Yao *et al*. (2018) to create the phylogeny seen in Fig. S 1.

| Species | Gene_ID | Gene_name |
| --- | --- | --- |
| *Marchantia polymorpha* | Mapoly0052s0098.1 | MapoPM1 |
| *Marchantia polymorpha* | Mapoly0022s0083.1 | MapoPM2 |
| *Marchantia polymorpha* | Mapoly0022s0082.1 | MapoPM3 |
| *Marchantia polymorpha* | Mapoly0022s0081.1 | MapoPM4 |
| *Marchantia polymorpha* | Mapoly0044s0120.1 | MapoPM5 |
| *Marchantia polymorpha* | Mapoly0022s0022.1 | MapoPM6 |
| *Sphagnum fallax* | Sphfalx0018s0035.1 | SpfaPM1 |
| *Sphagnum fallax* | Sphfalx0006s0197.1 | SpfaPM2 |
| *Sphagnum fallax* | Sphfalx0176s0017.1 | SpfaPM3 |
| *Sphagnum fallax* | Sphfalx0044s0087.1 | SpfaPM4 |
| *Sphagnum fallax* | Sphfalx0106s0009.1 | SpfaPM5 |
| *Sphagnum fallax* | Sphfalx0027s0209.1 | SpfaPM6 |
| *Sphagnum fallax* | Sphfalx0003s0261.1 | SpfaPM7 |
| *Sphagnum fallax* | Sphfalx0142s0026.1 | SpfaPM8 |
| *Sphagnum fallax* | Sphfalx0205s0031.1 | SpfaPM9 |
| *Sphagnum fallax* | Sphfalx0019s0102.1 | SpfaPM10 |
| *Sphagnum fallax* | Sphfalx0002s0395.1 | SpfaPM11 |
| *Sphagnum fallax* | Sphfalx0006s0346.1 | SpfaPM12 |
| *Sphagnum fallax* | Sphfalx0007s0073.1 | SpfaPM13 |
| *Zostera marina* | Zosma440g00070.1 | ZomaPM1 |
| *Zostera marina* | Zosma88g00500.1 | ZomaPM2 |
| *Zostera marina* | Zosma44g01500.1 | ZomaPM3 |
| *Setaria italica* | Seita.2G115100.1 | SeitPM1 |
| *Setaria italica* | Seita.9G002100.1 | SeitPM2 |
| *Setaria italica* | Seita.3G250100.1 | SeitPM3 |
| *Setaria italica* | Seita.5G284700.1 | SeitPM4 |
| *Setaria italica* | Seita.5G000700.1 | SeitPM5 |
| *Setaria italica* | Seita.9G230400.1 | SeitPM6 |
| *Setaria italica* | Seita.7G125100.1 | SeitPM7 |
| *Setaria italica* | Seita.8G228600.1 | SeitPM8 |
| *Brachypodium distachyon* | Bradi1g00600.1 | BrdiPM1 |
| *Brachypodium distachyon* | Bradi2g47520.2 | BrdiPM2 |
| *Brachypodium distachyon* | Bradi2g25960.1 | BrdiPM3 |
| *Brachypodium distachyon* | Bradi2g08680.1 | BrdiPM4 |
| *Solanum lycopersicum* | Solyc05g053160.2.1 | SolyPM1 |
| *Solanum lycopersicum* | Solyc09g090800.1.1 | SolyPM2 |
| *Solanum lycopersicum* | Solyc02g071310.2.1 | SolyPM3 |
| *Solanum lycopersicum* | Solyc01g080290.2.1 | SolyPM4 |
| *Solanum lycopersicum* | Solyc02g085770.2.1 | SolyPM5 |
| *Eucalyptus grandis* | Eucgr.H00118.1 | EugrPM1 |
| *Eucalyptus grandis* | Eucgr.H02898.1 | EugrPM2 |
| *Eucalyptus grandis* | Eucgr.D00344.1 | EugrPM3 |
| *Eucalyptus grandis* | Eucgr.H03634.1 | EugrPM4 |
| *Carica papaya* | Capa_evm.model.supercontig_16.51 | CapaPM1 |
| *Carica papaya* | Capa_evm.model.supercontig_179.19 | CapaPM2 |
| *Carica papaya* | Capa_evm.model.supercontig_17.171 | CapaPM3 |
| *Carica papaya* | Capa_evm.model.supercontig_179.15 | CapaPM4 |
| *Carica papaya* | Capa_evm.model.supercontig_59.22 | CapaPM5 |
| *Capsella rubella* | Carubv10002111m | CaruPM1 |
| *Capsella rubella* | Carubv10011510m | CaruPM2 |
| *Capsella rubella* | Carubv10027229m | CaruPM3 |
| *Capsella rubella* | Carubv10012389m | CaruPM4 |
| *Glycine max* | Glyma.20G184600.1 | GlmaPM1 |
| *Glycine max* | Glyma.10G205900.1 | GlmaPM2 |
| *Glycine max* | Glyma.09G085600.1 | GlmaPM3 |
| *Glycine max* | Glyma.15G195600.1 | GlmaPM4 |
| *Glycine max* | Glyma.08G121700.1 | GlmaPM5 |
| *Glycine max* | Glyma.05G164300.1 | GlmaPM6 |
| *Glycine max* | Glyma.07G255100.1 | GlmaPM7 |
| *Glycine max* | Glyma.17G019200.1 | GlmaPM8 |
| *Medicago truncatula* | Medtr1g094035.1 | MetrPM1 |
| *Medicago truncatula* | Medtr2g049340.1 | MetrPM2 |
| *Medicago truncatula* | Medtr6g068860.1 | MetrPM3 |
| *Medicago truncatula* | Medtr4g117600.1 | MetrPM4 |
| *Azolla filiculoides* | Azfi_s0001.g000130 | AzfiPM1 |
| *Azolla filiculoides* | Azfi_s0050.g031077 | AzfiPM2 |
| *Azolla filiculoides* | Azfi_s0185.g056678 | AzfiPM3 |
| *Azolla filiculoides* | Azfi_s0001.g000342 | AzfiPM4 |
| *Azolla filiculoides* | Azfi_s0083.g038896 | AzfiPM5 |
| *Azolla filiculoides* | Azfi_s0003.g007412 | AzfiPM6 |
| *Salvinia cucullata* | Sacu_v1.1_s0019.g007798 | SacuPM1 |
| *Salvinia cucullata* | Sacu_v1.1_s0101.g019776 | SacuPM2 |
| *Salvinia cucullata* | Sacu_v1.1_s0084.g018226 | SacuPM3 |
| *Anthoceros agrestis* | AagrOXF_evm.model.utg000057l.54.1<br>EVM AagrOXF_evm.TU.utg000057l.54<br>utg000057l:248554-249384(+) | AnagPM1 |
| *Anthoceros agrestis* | AagrOXF_evm.model.utg000057l.14.1<br>EVM AagrOXF_evm.TU.utg000057l.14<br>utg000057l:74404-75215(-) | AnagPM2 |
| *Anthoceros agrestis* | AagrOXF_evm.model.utg000004l.46.1 .<br>AagrOXF_evm.TU.utg000004l.46<br>utg000004l:259567-260346(+) | AnagPM3 |
| *Anthoceros agrestis* | AagrOXF_evm.model.utg000004l.47.1 .<br>AagrOXF_evm.TU.utg000004l.47<br>utg000004l:261096-262148(+) | AnagPM4 |
| *Anthoceros agrestis* | AagrOXF_evm.model.utg000004l.47.2 .<br>AagrOXF_evm.TU.utg000004l.47<br>utg000004l:261096-262086(+) | AnagPM5 |
| *Anthoceros agrestis* | AagrOXF_evm.model.utg000004l.47.3<br>AagrOXF_evm.TU.utg000004l.47<br>utg000004l:261096-261962(+) | AnagPM6 |
| *Anthoceros agrestis* | AagrOXF_evm.model.utg000095l.55.1<br>EVM AagrOXF_evm.TU.utg000095l.55<br>utg000095l:269706-271270(+) | AnagPM7 |
| *Physcomitrium (Physcomitrella)*<br>*patens* | Pp3c4_30620V3.1 | PhpaPM1 |
| *Physcomitrium (Physcomitrella)*<br>*patens* | Pp3c1_14380V3.1 | PhpaPM2 |
| *Physcomitrium (Physcomitrella)*<br>*patens* | Pp3c2_25930V3.1 | PhpaPM3 |
| *Physcomitrium (Physcomitrella)*<br>*patens* | Pp3c11_14870V3.1 | PhpaPM4 |
| *Physcomitrium (Physcomitrella)*<br>*patens* | Pp3c11_14880V3.1 | PhpaPM5 |
| *Physcomitrium (Physcomitrella)*<br>*patens* | Pp3c18_17550V3.1 | PhpaPM6 |
| *Physcomitrium (Physcomitrella)*<br>*patens* | Pp3c4_3450V3.1 | PhpaPM7 |
| *Physcomitrium (Physcomitrella)*<br>*patens* | Pp3c7_11540V3.1 | PhpaPM8 |
| *Physcomitrium (Physcomitrella)*<br>*patens* | Pp3c12_25110V3.1 | PhpaPM9 |
| *Physcomitrium (Physcomitrella)*<br>*patens* | Pp3c4_3440V3.1 | PhpaPM10 |
| *Physcomitrium (Physcomitrella)*<br>*patens* | Pp3c20_13220V3.1 | PhpaPM11 |
| *Physcomitrium (Physcomitrella)*<br>*patens* | Pp3c4_21960V3.1 | PhpaPM12 |
| *Selaginella moellendorffii* | Semo439591 | SemoPM1 |
| *Selaginella moellendorffii* | Semo416876 | SemoPM2 |
| *Selaginella moellendorffii* | Semo444324 | SemoPM3 |
| *Selaginella moellendorffii* | Semo37585 | SemoPM4 |
| *Selaginella moellendorffii* | Semo437726 | SemoPM5 |
| *Selaginella moellendorffii* | Semo432371 | SemoPM6 |
| *Ginkgo biloba* | Gb_19410 | GibiPM1 |
| *Ginkgo biloba* | Gb_35411 | GibiPM2 |
| *Ginkgo biloba* | Gb_35398 | GibiPM3 |
| *Amborella trichopoda* | AmTr_v1.0_scaffold00039.123 | AmtrPM1 |
| *Amborella trichopoda* | AmTr_v1.0_scaffold00005.58 | AmtrPM2 |
| *Amborella trichopoda* | AmTr_v1.0_scaffold00025.112 | AmtrPM3 |
| *Aquilegia coerulea* | Aqcoe5G012300.1 | AqcoPM1 |
| *Aquilegia coerulea* | Aqcoe3G398700.1 | AqcoPM2 |
| *Aquilegia coerulea* | Aqcoe1G207600.1 | AqcoPM3 |
| *Aquilegia coerulea* | Aqcoe2G327400.1 | AqcoPM4 |
| *Arabidopsis thaliana* | AT1G04560.1 | AtPM19L1 |
| *Arabidopsis thaliana* | AT5G46530.1 | AtPM19L2 |
| *Arabidopsis thaliana* | AT1G29520.1 | AtPM19L3 |
| *Arabidopsis thaliana* | AT5G18970.1 | AtPM19L4 |
| *Musa acuminata* | GSMUA_Achr8P12720_1 | MuacPM1 |
| *Musa acuminata* | GSMUA_Achr11P02400_1 | MuacPM2 |
| *Musa acuminata* | GSMUA_Achr2P20160_1 | MuacPM3 |
| *Musa acuminata* | GSMUA_Achr4P13350_1 | MuacPM4 |
| *Musa acuminata* | GSMUA_Achr8P06030_1 | MuacPM5 |
| *Oryza sativa* | LOC_Os05g31670.1 | OsPM1 |
| *Oryza sativa* | LOC_Os07g24000.1 | OsPM2 |
| *Oryza sativa* | LOC_Os02g49860.1 | OsPM3 |
| *Oryza sativa* | LOC_Os01g50440.1 | OsPM4 |
| *Oryza sativa* | LOC_Os01g14530.1 | OsPM5 |
| *Oryza sativa* | LOC_Os05g33220.1 | OsPM6 |
| *Oryza sativa* | LOC_Os10g32720.1 | OsPM7 |

**Table S3.** Yeast Strains.

| Strain | Description | Source |
| --- | --- | --- |
| JRY379 | MATa his3, leu2, trp1, ura3, suc2 | Prof. Per Ljungdahl,<br>Ludwig institute for<br>Cancer Research,<br>Stockholm, Sweden |
| PLY240 | MATa his3, leu2, trp1, ura3, suc2, trk1D51<br>trk2D50::lox-kanMX-lox | Prof. Per Ljungdahl,<br>Ludwig institute for<br>Cancer Research,<br>Stockholm, Sweden |
| W303-1A | MATa, leu2, ura3, trp1, his3, ade2, can1-100, GAL,<br>SUC2, mal0 | Prof. Roberto A.<br>Gaxiola, Arizona<br>State University, USA |
| YSH642 | MATa, leu2-3/112 ura3-1 trp1-1 his3-11/15 ade2-1<br>can1-100 GAL SUC2, mal0, gpd1D::TRP1 gpd2D::URA3 | Prof. Stefan<br>Hohmann,<br>Chalmers University<br>of Technology<br>Göteborg, Sweden |
| L5709 | MATa, leu2, ura3, trp1, his3, ade2, can1-100, GAL,<br>SUC2, mal0, ena1::HIS3 | Prof. Roberto A.<br>Gaxiola (Arizona<br>State University,<br>USA) |
| TM100 | MATa, ura3, leu2, trp1 | Prof T. Maeda,<br>Cancer Institute,<br>Tokyo, Japan |
| TM310 | MATa ura3 leu2 trp1 his3 ssk2::URA3 ssk22::LEU2<br>sho1::TRP1 | Prof T. Maeda,<br>Cancer Institute,<br>Tokyo, Japan |
| NMY51 | MATa, his3, trp1, leu2, ade2, LYS2::(lexAop)4-HIS3<br>ura3::(lexAop)8-lacZ ade2::(lexAop)8-ADE2 GAL | Dualsystems Biotech<br>AG |

**Table S4.** Plasmids used in this work

| Plasmid | Features | Use | Source |
| --- | --- | --- | --- |
| *Complementation assays* |  |  |  |
| pCambia 1302 | CaMV35S:GFP-His | Plant transformation | Prof. Richard Jefferson, Cambia, Australia |
| pCaMV35S:PM19L1-GFP | pCaMV35S:PM19L1-GFP | Plant transformation | This study |
| pPM19L1:PM19L1 | Native PM19L1 promoter and gene | Plant transformation | This study |
| pPR97 | UidA | Plant transformation | Prof. P. Ratet, CNRS, France |
| pPM19L1:UidA | Promoter PM19L1:UidA | Plant transformation | This study |
| pYES2 | 2 μm URA3 PGal- CYC1 TT | Expression in yeast | Invitrogen |
| pPMRA1 | PM19L1 in pYES2 | Complementation of<br>PLY240 (*trk1/trk2*)<br>RGY85 (*ena1*) | This study |
| pPMRA2 | LEU2 replacement of URA3 in pYES2 | Expression in yeast | This study |
| pPMRA3 | PM19L1 in pPMRA2 | Complementation of<br>YSH642 (*gdp1/gdp2*) | This study |
| pPMRA4 | HIS3 replacement of | Expression in yeast | This study |
|  | URA3 in pYES2 |  |  |
| pPMRA5 | PM19L1 in pPMRA4 | Complementation of TM310<br>(*ssk2/ssk22/sho1*) | This study |
| pPMRA6 | PM19L1Δ in pPMRA4 | Complementation of TM310<br>(*ssk2/ssk22/sho1*) | This study |
| pPMRA7 | *Kan(R)* replacement of URA3 in pYES2 | Expression in yeast | This study |
| pPMRA8 | MKK1 in pPMRA7 | Enhance complementation of TM310<br>(*ssk2/ssk22/sho1*) | This study |
| pPMRA9 | MKK2 in pPMRA6 | Enhance complementation of TM310<br>(*ssk2/ssk22/sho1*) | This study |
| pPMRA10 | MKK3 in pPMRA6 | Enhance complementation of TM310<br>(*ssk2/ssk22/sho1*) | This study |
| pPMRA11 | Mutated PM19L1 (79W-G), in pPMRA5 | complementation of TM310<br>(*ssk2/ssk22/sho1*) | This study |
| pPMRA12 | Mutated PM19L1 (80R-G), in pPMRA5 | complementation of TM310<br>(*ssk2/ssk22/sho1*) | This study |
| pPMRA13 | Mutated PM19L1, (82D-G) in pPMRA5 | complementation of TM310<br>(*ssk2/ssk22/sho1*) | This study |
| pPMRA14 | Tetraspanin in pPMRA4 | Complementation of TM310<br>(*ssk2/ssk22/sho1*) | This study |
| *Split ubiquitin assays* |  |  |  |
| pOst1-Nubl | 2 μm, TRP | 'Positive' prey vector | Iyer *et al.*, 2005 |
| pOst1-NubG | 2 μm, TRP | 'Negative' prey vector | Iyer *et al.*, 2005 |
| pPR3-C | TRP1 pADH1 - HANubG | Prey vector | Snider *et al.*, 2010 |
| pAMBV | LEU2 pADH1 - Cub-LexA-VP16 tag | Bait vector | Snider *et al.*, 2010 |
| pPMRA15 | LEU2 pADH1 - PM19L1 Cub-LexA-VP16 tag | Bait vector | This study |
| pPMRA16 | LEU2 pADH1 - PM19L1Δ Cub-LexA-VP16 tag | Bait vector | This study |
| pPMRA17 | LEU2 pADH1 - PM19L1(**79W-G**) Cub-LexA-VP16 tag | Bait vector | This study |
| pPMRA18 | LEU2 pADH1- PM19L1(**80R-G**) Cub-LexA-VP16 tag | Bait vector | This study |
| pPMRA19 | LEU2 pADH1- PM19L1(**81N-G**) Cub- | Bait vector | This study |
|  | LexA-VP16 tag |  |  |
| pPMRA20 | LEU2 pADH1-PM19L1(82D-G) Cub-LexA-VP16 tag | Bait vector | This study |
| pPMRA21 | TRP1 pADH1 -MKK1-HANubG | Prey vector | This study |
| pPMRA22 | TRP1 pADH1 -MKK2-HANubG | Prey vector | This study |
| pPMRA23 | TRP1 pADH1 -MKK3-HANubG | Prey vector | This study |
| *Fusion protein production* |  |  |  |
| pMAL-C2 | Maltose binding protein | Fusion protein<br>production | New England Biolab |
| pMAL-MKK1 | MBP-MKK1 fusion | Recombinant MKK1 | Huang et al, 2000 |
| pMAL-MKK2 | MBP-MKK2 fusion | Recombinant MKK2 | This study |
| pMAL-MKK3 | MBP-MKK3 fusion | Recombinant MKK3 | This study |

**Table S5.**
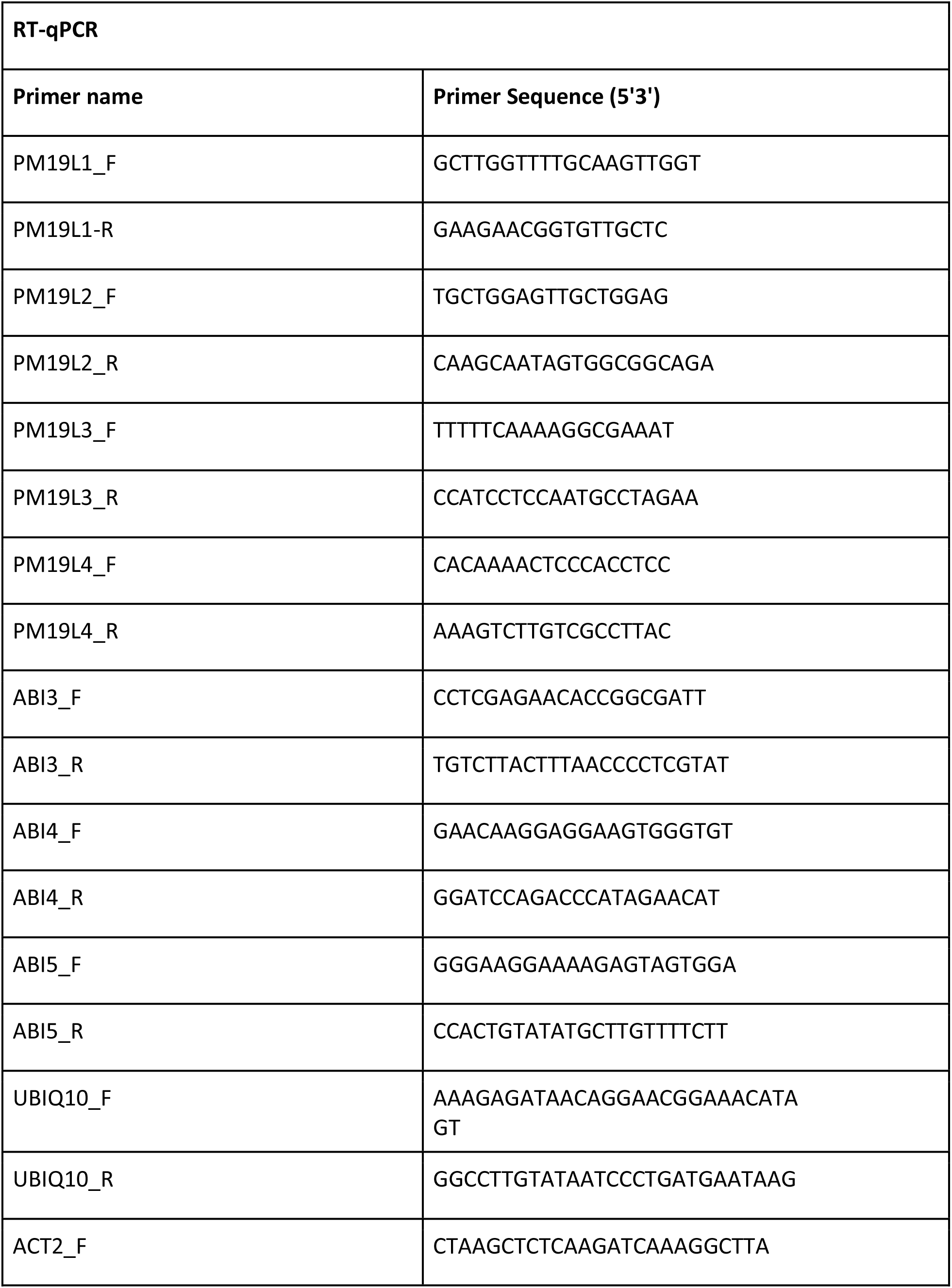
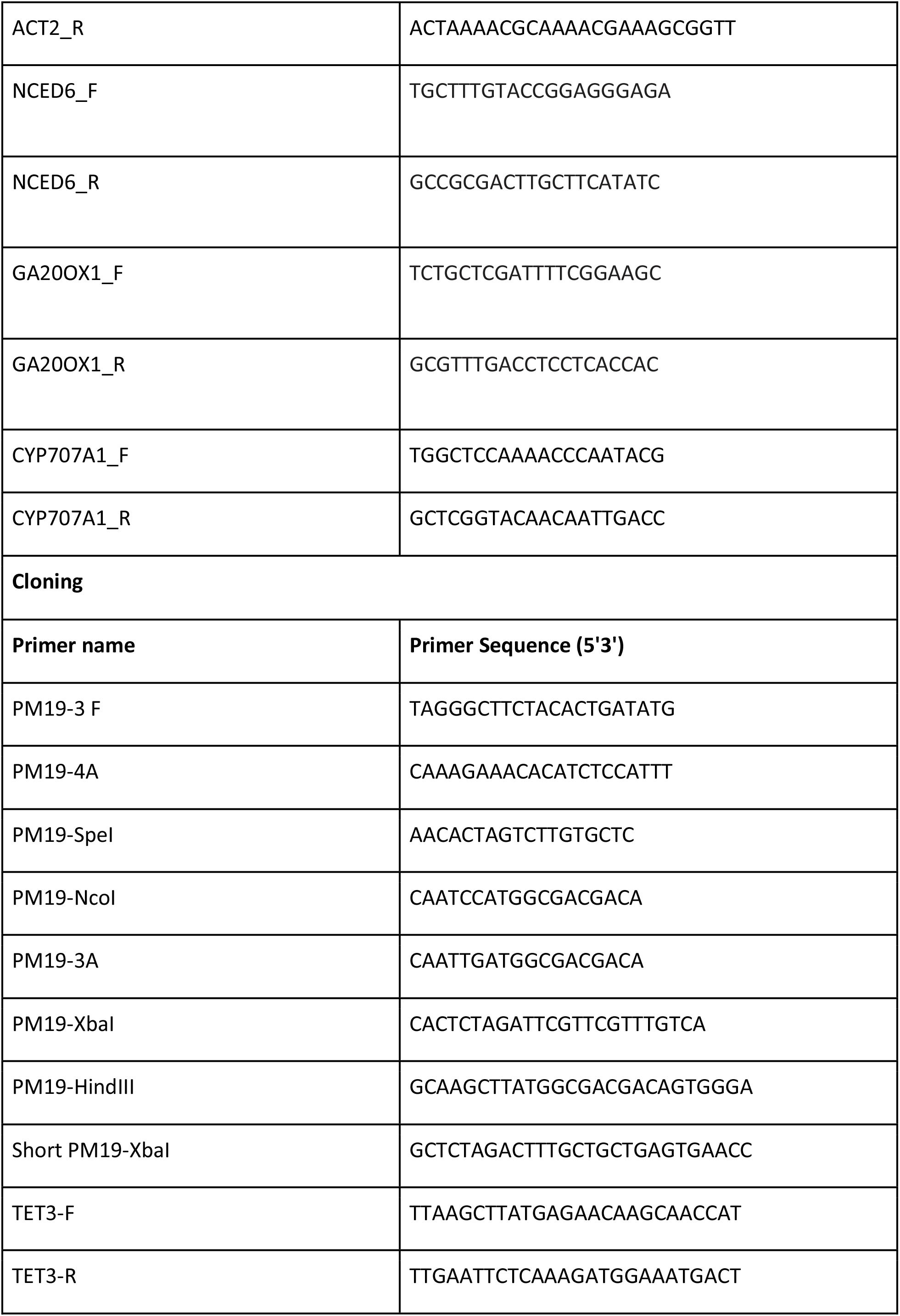
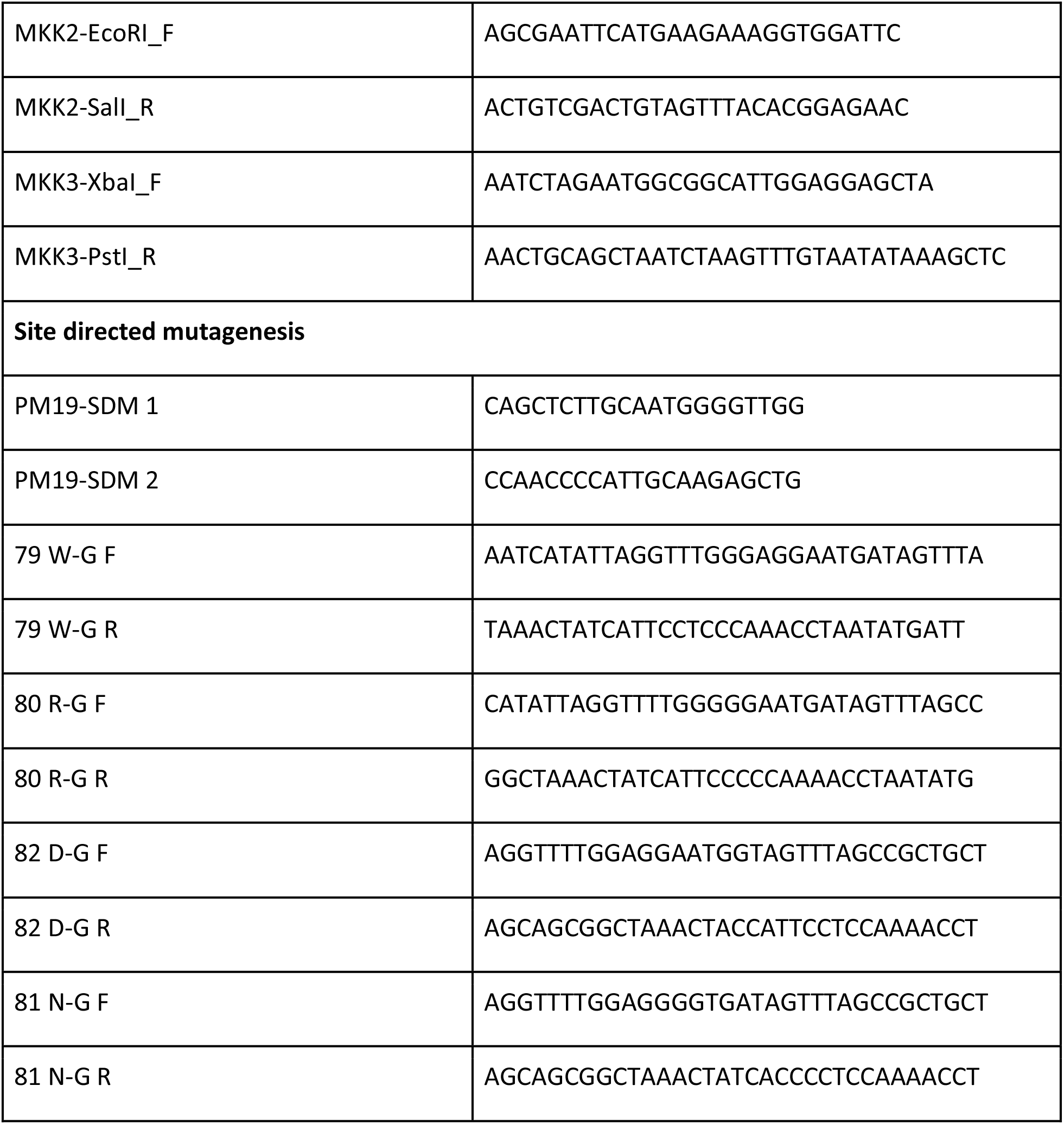
DNA primers used in this study.

